# Molecular underpinnings of ssDNA specificity by Rep HUH-endonucleases and implications for HUH-tag multiplexing and engineering

**DOI:** 10.1101/2020.09.01.278671

**Authors:** KJ Tompkins, M Houtti, LA Litzau, EJ Aird, BA Everett, AT Nelson, L Pornschloegl, LK Limón-Swanson, RL Evans, K Evans, K Shi, H Aihara, WR Gordon

## Abstract

Replication initiator proteins (Reps) from the HUH-endonuclease superfamily process specific single-stranded DNA (ssDNA) sequences to initiate rolling circle/hairpin replication in viruses, such as crop ravaging geminiviruses and human disease causing parvoviruses. In biotechnology contexts, Reps are the basis for HUH-tag bioconjugation and a critical adeno-associated virus genome integration tool. We solved the first co-crystal structures of Reps complexed to ssDNA, revealing a key motif for conferring sequence specificity and for anchoring a bent DNA architecture. In combination, we developed a deep sequencing cleavage assay, termed HUH-seq, to interrogate subtleties in Rep specificity and demonstrate how differences can be exploited for multiplexed HUH-tagging. Together, our insights allowed engineering of only four amino acids in a Rep chimera to predictably alter sequence specificity. These results have important implications for modulating viral infections, developing Rep-based genomic integration tools, and enabling massively parallel HUH-tag barcoding and bioconjugation applications.

## Introduction

HUH-endonucleases are diverse enzymes utilizing common single-stranded sDNA (ssDNA) processing mechanisms that break and join DNA to facilitate fundamental biological processes such as rolling circle replication (RCR), rolling hairpin replication (RHR), bacterial conjugation, DNA transposition, and DNA integration into host genomes ^1–3^. At the heart of DNA processing of all HUH-endonucleases is a structurally defined catalytic nickase domain that first recognizes a specific sequence/structure of DNA; nicks ssDNA at a “*nic* site” to yield a sequestered 5’ end that remains covalently bound to the HUH endonuclease and a free 3’OH that can be used as a primer for DNA replication; and finally, facilitates a strand transfer reaction to resolve the covalent intermediate ^1^. The two major classes of HUH-endonucleases are replication initiator proteins (Reps) involved in RCR and RHR and relaxases involved in bacterial conjugation of plasmids, although HUH-endonucleases are also involved in DNA transposition ^1^.

The covalent phosphotyrosine intermediate has recently been exploited for biotechnology applications. “HUH-tag” fusion proteins are emerging as a versatile bioconjugation platform to covalently link proteins to DNA, combining the diverse functionality of proteins with the programmability of DNA ^4^. HUH-tag applications have permeated into technologies such as DNA origami scaffolded protein assembly ^5–8^, receptor-specific cell targeting by adeno-associated virus ^9^, aptamer-based sandwich detection ^10^, directed nanoparticle drug-delivery via DNA aptamers ^11^, and CRISPR-Cas9 genome engineering ^12,13^, mainly due to their ability to form robust covalent adducts under physiologic conditions. Rather than relying on expensive nucleic acid modifications such as the SNAP-tag ^14^, CLIP-tag ^15^, and Halo-tag ^16^ systems, HUH-tags rely on an inherent ssDNA binding moiety that promotes the catalysis of a transesterification reaction resulting in a stable phosphotyrosine adduct ^1^.

Understanding the molecular basis of DNA recognition by HUH-endonucleases could provide much needed solutions for bacterial antibiotic resistance resulting from HUH-endonuclease mediated horizontal gene transfer ^17^, as well as in the prevention or treatment of HUH-endonuclease mediated viral infections, such as geminivirus infections of plants that ravage the agricultural crop industry ^18,19^ and parvovirus B19 infections of humans ^20^ that are associated with a range of autoimmune diseases ^21,22^. Moreover, the ability to rationally engineer HUH-endonucleases to recognize a desired DNA sequence has huge potential in genome engineering ^23^ and DNA delivery applications as well as in expanding the multiplexibility of HUH-tagging to meet the demand of the recent explosion of DNA-barcoding applications ^24–27^.

However, while several structures of relaxase HUH-endonucleases in complex with their cognate DNA target sequences have been reported ^17,28–30^, there are no structures of viral Rep HUH-endonucleases in complex with ssDNA comprising the target sequence at the origin of replication (*ori*). Despite structurally superimposable active sites and a common overall core structure ^31^, there are several structural elements of the larger relaxase proteins that do not exist in Reps, such as extensions of the C-terminus and internal loops with respect to Reps. These structures form extensive contacts with the target DNA, thus underscoring potential differences in DNA recognition mechanisms between Reps and relaxases ^32^.

In this study, we determined the structural basis for ssDNA recognition by viral Rep HUH-endonucleases by solving two Rep-ssDNA co-crystal structures and identified an ssDNA “bridging” motif largely responsible for DNA recognition. To further interrogate the ssDNA specificity of Reps, we developed HUH-seq, a high-throughput, next generation sequencing (NGS)-based DNA cleavage assay that we used to define ssDNA recognition profiles of a panel of ten Reps using a ssDNA library containing 16,384 different target sequences. Despite the high similarity of cognate nonanucleotide *ori* sequences and the promiscuous nature of Rep ssDNA recognition, HUH-seq analysis surprisingly revealed many examples of orthogonal adduct formation between Reps from different viral families with little or no cross-reactivity. Finally, we rationally engineered a chimeric Rep by swapping a few amino acids of the ssDNA “bridging” motif of one Rep into the backbone of a related Rep, predictably modulating ssDNA sequence specificity.

## Results

### REP HUH-ENDONUCLEASE CO-CRYSTAL STRUCTURES

To uncover the ssDNA recognition mechanism of Reps and identify potential motifs that might confer sequence specificity, we solved the first high resolution crystal structures of two distinct Rep HUH-endonuclease domains, from porcine circovirus 2 (PCV2) and wheat dwarf virus (WDV), bound to ssDNA encoding minimal sequences comprising the respective *ori* sequences. The pre-cleavage state was captured by mutating the catalytic tyrosine to phenylalanine. We present structures of two 10-mer bound structures of inactive PCV2^Y96F^ (1.93 Å resolution) and WDV^Y106F^ (1.80 Å resolution), and one 8-mer bound WDV^Y106F^ (2.61 Å resolution) structure (Table 1). The three structures are in complex with the divalent cofactor manganese, and the catalytic tyrosine (though a Phe mutant in the structures) is positioned for nucleophilic attack of the scissile phosphate (Fig. 1a and 1b, Supp. Note 1). PCV2 and WDV share a highly similar ssDNA binding interface, albeit with several distinct features.

**Table 1:** Data collection and refinement statistics.

|  | PCV<sup>Y96F</sup> + 10mer<sup>a</sup><br>(PDB 6WDZ) | WDV<sup>Y106F</sup> + 10mer<sup>a</sup><br>(PDB 6WE0) | WDV<sup>Y106F</sup> + 8mer<sup>a</sup><br>(PDB 6WE1) |
| --- | --- | --- | --- |
| **Data Collection** |  |  |  |
| Wavelength (Å) | 0.979 | 0.979 | 1.542 |
| Resolution range (Å) | 86.26 - 1.93 (1.99 - 1.93)<sup>b</sup> | 41.34 - 1.8 (1.864 - 1.8) | 28.55 - 2.61 (2.71 - 2.61) |
| Space group | *P* 6<sub>4</sub> | *P* 2<sub>1</sub> 2<sub>1</sub> 2<sub>1</sub> | *P* 4<sub>1</sub> |
| Unit cell (Å) | *a* = *b* = 99.61<br>*c* = 73.72<br>90° 90° 120° | *a* = 45.57<br>*b* = 50.01 *c* = 73.44<br>90° 90° 90° | *a* = *b* = 50.63<br>*c* = 241.98<br>90° 90° 90° |
| Unique reflections | 30660 (3087) | 15366 (6674) | 15929 (1507) |
| Completeness (%) | 97.72 (97.87) | 95.23 (98.54) | 86.62 (82.15) |
| Wilson B-factor | 28.81 | 26.51 | 52.23 |
| Mean I/σ | 9.46 (1.67) | 14.17 (2.05) | 7.70 (2.27) |
| CC1/2 | 0.995 (0.520) | 0.997 (0.571) | 0.987 (0.702) |
| CC* | 0.999 (0.827) | 0.999 (0.862) | 0.997 (0.908) |
| R-meas | 0.130 (1.083) | 0.0836 (0.963) | 0.112 (0.609) |
| R-pim | 0.064 (0.528) | 0.037 (0.436) | 0.071 (0.378) |
| **Refinement** |  |  |  |
| Reflections used in refinement | 30657 (3085) | 15364 (1550) | 15917 (1505) |
| Reflections used for R-free | 1996 (206) | 818 (92) | 807 (66) |
| R-work | 0.245 (0.366) | 0.173 (0.220) | 0.167 (0.264) |
| R-free | 0.301 (0.406) | 0.224 (0.313) | 0.236 (0.326) |
| Number of non-hydrogen atoms | 3290 | 1169 | 4321 |
| macromolecules | 3131 | 1112 | 4278 |
| ligands | 3 | 4 | 6 |
| solvent | 156 | 53 | 37 |
| Protein residues | 312 | 119 | 471 |
| RMS(bonds) (Å) | 0.010 | 0.006 | 0.008 |
| RMS(angles) (deg.) | 1.14 | 1.09 | 1.06 |
| Ramachandran favored (%) | 92.74 | 98.29 | 95.81 |
| Ramachandran allowed (%) | 4.95 | 1.71 | 4.19 |
| Ramachandran outliers (%) | 2.31 | 0 | 0 |
| Rotamer outliers (%) | 3.45 | 0 | 0.27 |
| Clashscore | 14.54 | 0 | 4.84 |
| Average B-factor | 38.02 | 30.37 | 47.84 |
| macromolecules | 38.11 | 29.89 | 47.84 |
| ligands | 29.34 | 34.76 | 44.29 |
| solvent | 36.55 | 40.15 | 48.76 |

a = Data are from one crystal
b = values in parentheses are for highest resolution shell

**Fig. 1:**
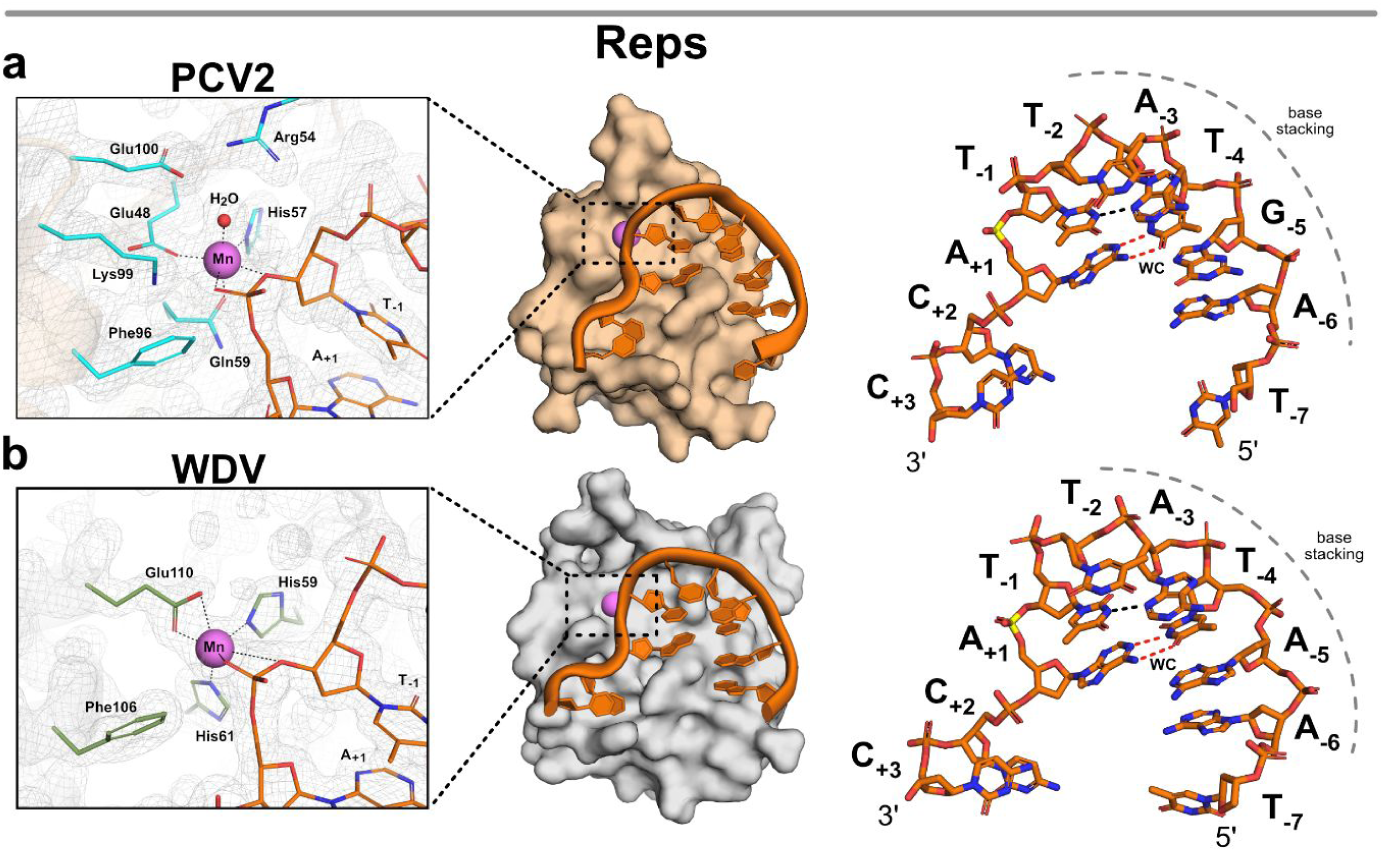
PCV and WDV Rep co-crystal structures complexed with ssDNA target sequence. **a**, Surface representation of PCV2^Y96F^ colored in beige and **b**, WDV^Y106F^ colored in gray bound to manganese as a sphere in magenta and DNA 10-mers as sticks colored orange by element. PCV2^Y96F^ is bound to 10-mer (5’-dTAGTATTACC-3’), and WDV^Y106F^ is bound to 10-mer (5’-dTAATATTACC-3’) both adopting a U-shaped conformation. Nucleotides are labeled as single letter abbreviations and positions, indicated as subscripts, relative to the scissile phosphate in yellow. A dashed gray curve indicates the base stacking chain that occurs between positions −6 through −2. Intramolecular Watson-Crick (WC) base pairing between A_+1_ and T_-4_ are indicated by red dashed lines as well as a non-canonical hydrogen bond between T_-1_ and A_-3_ are indicated as a black dashed line. Active site side chains are indicated as sticks, PCV2^Y96F^ in cyan and WDV^Y106F^ in green by element. The PCV2^Y96F^ active site coordinates the manganese in an octahedral geometry using Glu48, His57, and Gln59 with a water and two oxygens of the scissile phosphate completing the coordination shown as black dashed lines. The WDV^Y106F^ active site coordinates the manganese in an octahedral geometry using Glu110, His59, and His61 with two oxygens of the scissile phosphate completing the coordination shown as black dashed lines. The active site is displayed within the 2Fo-Fc map mesh at σ = 2.

### REP DNA DOCKING INTERFACE CONFORMS ssDNA TO “U-SHAPED” ARCHITECTURE

Reps involved in RCR are known to cleave in the loop of a DNA hairpin harboring the cognate *ori* sequence (Supp. Fig 1). Strikingly, despite the absence of bases that make up the hairpin stem in the short target DNA oligos, the ssDNA is bent into a “U-shaped” architecture like one might expect in the context of the hairpin loop. The U-shaped DNA sits in a shallow channel on the surface of one face of the Rep protein with a distinct topological “nose” that juts out in the center of the U. The bent conformation of the ssDNA in the Rep structures is driven by both intermolecular interactions with the topological “nose” of the protein and by intramolecular Watson-Crick base pairing between T_-4_ and A_+1_ along with adjacent hydrogen bonding between N3 of T_-1_ and N3 of A_-3_ (Fig. 1a and 1b). Moreover, energetically favorable base stacking occurs between 5 nucleotides at positions −6 through −2. These intramolecular, conformation stabilizing interactions, along with protein-nucleotide interactions, promote the proper orientation needed for catalysis of the 5’ phosphate of the position +1 nucleotide.

To analyze the contacts between protein and DNA facilitating sequence-specific ssDNA recognition, we generated protein-nucleotide interaction maps utilizing the DNAproDB platform ^33,34^, which reports contacts within 4 Å between protein and ssDNA (Supp. Fig. 2). The relative positions of residues directly involved in forming the ssDNA docking interface, the catalytic tyrosine, and the divalent metal coordinating residues of the 10-mer bound Rep structures are depicted as a cartoon (Fig. 2a and 2b) and mapped onto a structure-based alignment of several Reps (Fig. 2c). The structural positioning of residues involved in protein-DNA contacts in the PCV2 and WDV are nearly conserved, while the residue identity is more divergent. A majority of the ssDNA docking interface is created by a stretch of 9-10 consecutive residues that partly correspond to the topological “nose” sticking up in the middle of the U, comprising an observed turn-*β*4-turn structural motif, which resides within a previously defined region termed the geminivirus recognition sequence (GRS) (Fig. 2c) ^35^. A second prominent cluster of protein-DNA contacts reside within Motif I, both of which were previously implicated in DNA binding ^35–37^.

**Fig. 2:**
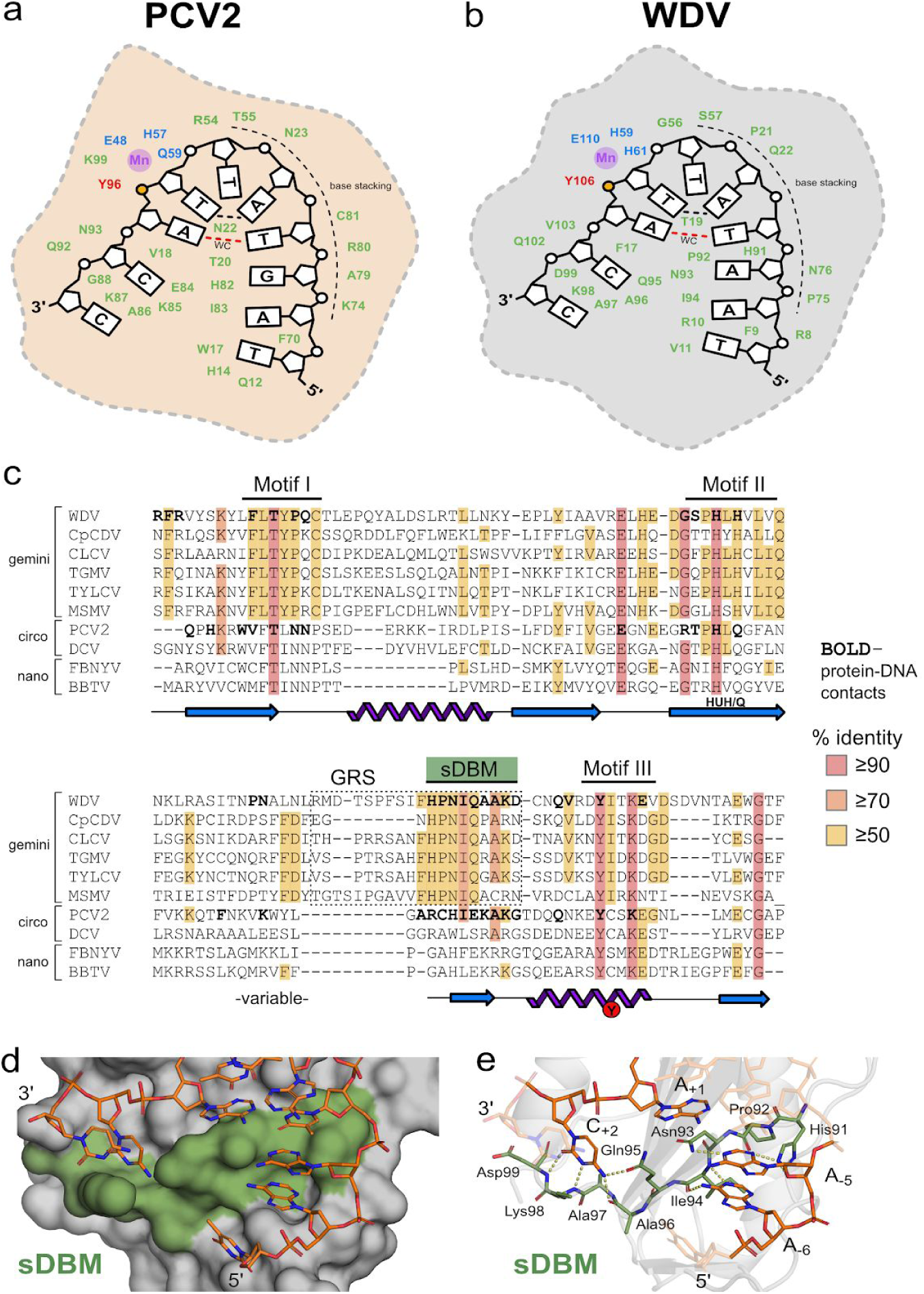
Cartoon depiction and structural alignment of specific Rep protein-DNA interactions. **a**, PCV2 and **b**, WDV Rep structures depicted as 2D cartoons with relative positions of residues (green) involved in binding “U-shaped” ssDNA within 4 Å. The catalytic tyrosine 106 is indicated in red with the adjacent phosphate in yellow, the ion coordinating triad is indicated in blue, and the 2+ ion in purple. The single Watson-Crick (WC) base pair is indicated as a dashed red line, and the ssDNA intramolecular hydrogen bond is indicated as a black dashed line. **c**, Structural alignment of Reps using PROMALS3D including available PCV2 (PDB: 6WDZ), WDV (PDB: 6WE0), TYLCV (1L2M), and FBNYV (6H8O) structures ^38–40^ as templates with conserved residues highlighted - high or absolute conservation (≥90%) indicated in red; moderately conserved (≥70%) indicated in orange; low conservation (≥50%) indicated in yellow; and no conservation (<50%) indicated in white. Amino- and carboxy-terminal ends are trimmed to reflect only structured domains in crystal structures. Bolded residues indicate contacts within 4 Å of DNA 10-mers complexed with PCV2^Y96F^ (PDB: 6WDZ), and WDV^Y106F^ (PDB: 6WE0). Conserved Rep Motifs I/II/III are shown as well as the GRS motif within the dashed box for geminivirus Reps. The sDBM we have defined in this study is labelled and highlighted in green. The conserved secondary structural elements (β1-5 and α1-2) below the alignment sequences are approximately shown as 2D cartoons with labeled HUH/Q motif and catalytic tyrosine. **d**, Surface representation of WDV with sDBM highlighted in green bound to the 10-mer as sticks. **e**, Major polar interactions between WDV sDBM residues (green sticks) and bases of 10-mer (orange sticks) are shown as yellow dashes.

### DEFINING THE SINGLE-STRANDED DNA BRIDGING MOTIF (sDBM)

The consecutive stretch of 9-10 residues in the turn-*β*4-turn structural motif (‘ARCHIEKAKG’ for PCV2 and ‘HPNIQAAKD’ for WDV) has two critical functions in the structure. First, it acts as a “bridge” between 5’ and 3’ ends of the nonanucleotide sequence contacting positions −6, −5, +1, and +2. (Fig 2d and 2e). In combination with the intramolecular base pairing and hydrogen bonding of the ssDNA, this sequence of residues likely contributes to bending and stabilizing the ssDNA in the U-shaped conformation. In the WDV^Y106F^ + 10-mer structure, residues His91 and Asp93 in this “bridging” motif specifically contact the base of A_-5_ (Fig. 2d and 2e), whereas Arg79 and His81 in the PCV2 structure specifically contact the base of G_-5_. We hypothesize that these specific contacts play a major role in conferring specificity differences at the −5 position. Previously, this motif remained undefined across Rep classes because of divergence in sequence conservation, though this divergence may be a major impetus for ssDNA recognition. Further, this motif is located in the N-terminus in relaxases and is involved in ssDNA binding (Supp. Fig. 3a and 3b). With this, we term this turn-*β*-turn structural motif as the ‘single-stranded DNA Bridging Motif’ (sDBM), and suggest that it is the main binding moiety responsible for recognition and conformation priming of ssDNA by Rep HUH-endonucleases.

**Fig. 3:**
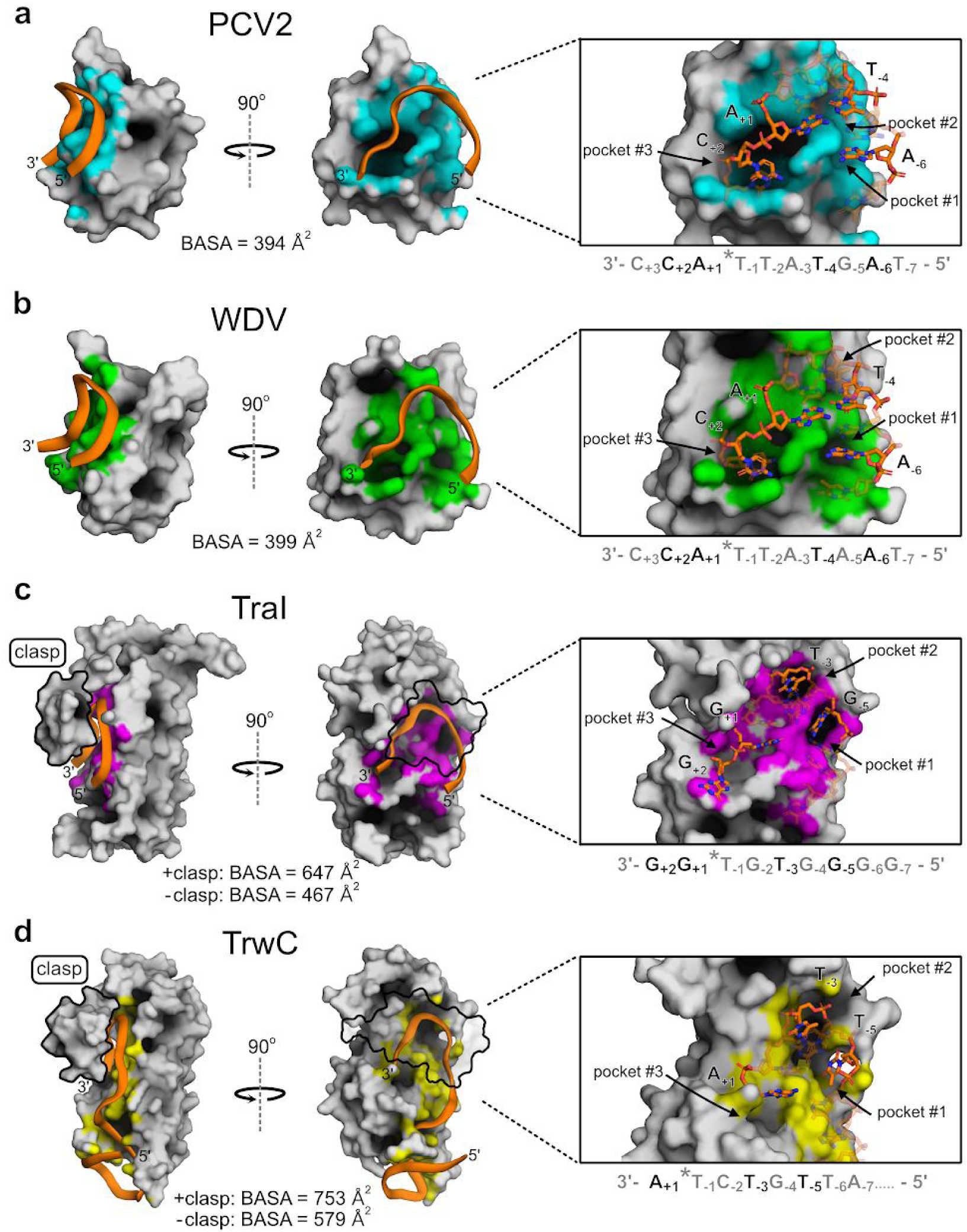
Structural comparison of ssDNA recognition by Reps and relaxases. **a-d**, PCV2 (6WDZ), WDV (6WE0), TraI36 (2A0I), TrwC (2CDM), are illustrated in gray surface display where DNA interactions within 4 Å are highlighted in cyan, green, magenta, and yellow, respectively. Bound single-stranded DNA is represented as a cartoon backbone in orange or as sticks with carbons and phosphates colored in orange. Nucleotides bound inside pockets are solid; other bound nucleotides are transparent. Relaxase “clasps” (TraI36 residues 231-271 and TrwC residues 237-262) are either solid or transparent and outlined in black. Total buried solvent accessible surface area (BASA; Å^2^) for ssDNA bound to the docking interface was calculated for each structure including values for with, or without, contribution from relaxase claps.

### REP VERSUS RELAXASE ssDNA RECOGNITION MECHANISM

Reps generally initiate replication of a large number of viruses and plasmids to copy their circular genomes while relaxases catalyze the transfer of one DNA strand of the plasmid genome to the recipient cell during plasmid conjugation ^1^; thus, relaxases are thought to recognize DNA with more specificity than Reps. Our structures provide insights at the molecular level into different modes of recognition between Reps and relaxases that should illuminate structural nuances of ssDNA recognition. The two available relaxase structures that are the most comparable to the Rep co-crystal structures are TraI (PDB ID: 2A0I) and TrwC (PDB ID: 2CDM), which are both complexed with ssDNA and have at least one nucleotide bound on the 3’ side of the *nic* site (Fig. 3c and 3d). Structurally, Reps and relaxases share a similar central 5-stranded antiparallel beta-sheet displaying the HUH motif, though the relaxases are circularly permuted with respect to the Reps such that the catalytic tyrosine is near the C-terminus of Reps and the N-terminus of relaxases ^31^. Relaxases have similar active sites and U-shaped ssDNA architectures to Reps ^28,30^. However, there are striking differences in how the two families of proteins recognize DNA. Aside from the most obvious difference of a larger size and a more extended DNA binding interface that includes binding a hairpin structure 5’ to the *nic* site of relaxase proteins, the most distinctive difference is that the relaxase structures contain a protein alpha-helical “clasp” that covers the bound DNA (Fig. 4). This clasp forms extensive contacts with the DNA, suggesting that it helps anchor the DNA to the protein. This is underscored by the fact that in the crystal structure of NES, the relaxase from staphylococcus aureus and which does not contain a “clasp”, the 3’ end of the DNA has very few contacts with the protein ^17^.

**Fig. 4:**
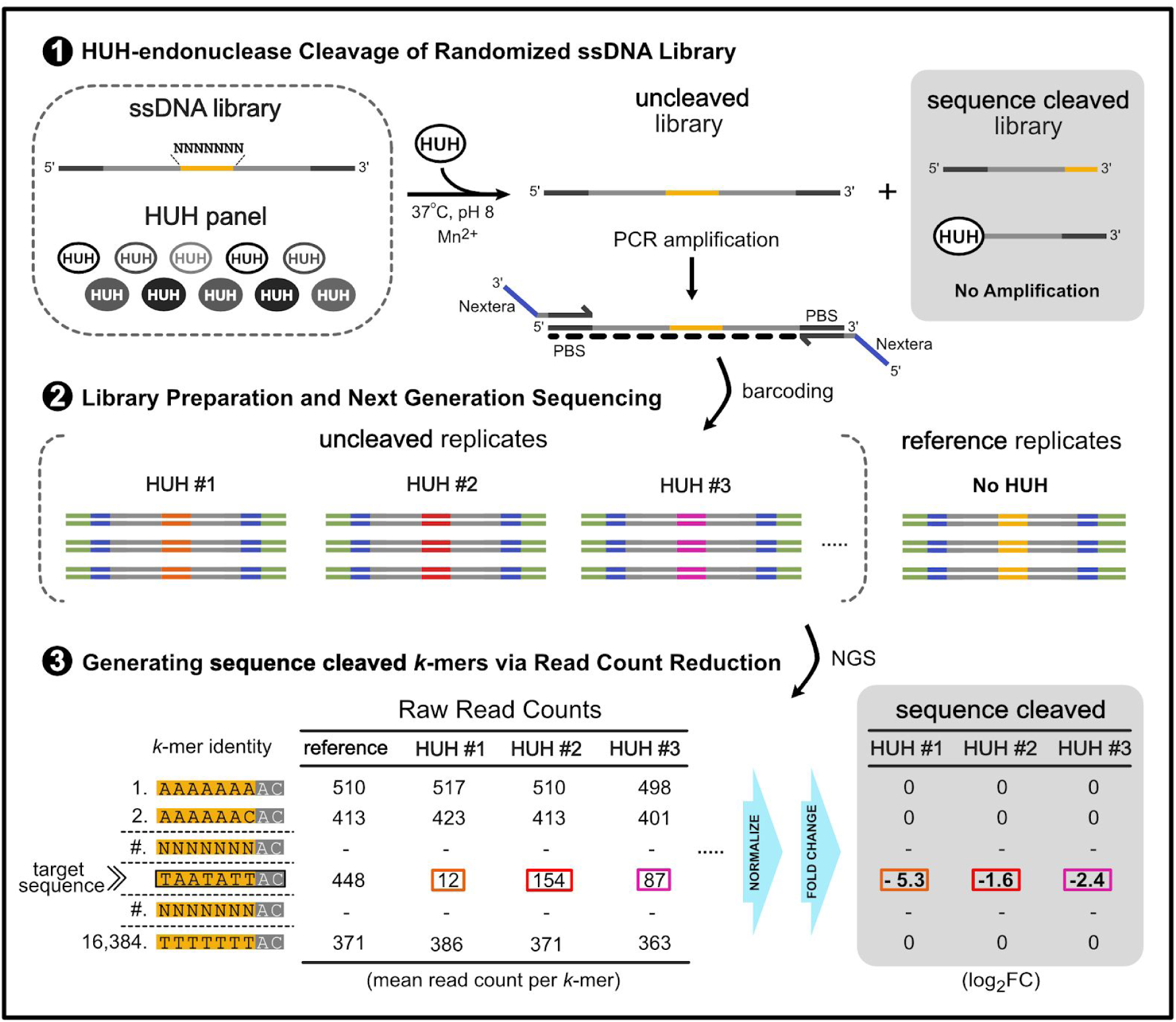
HUH-seq cleavage assay schematic for determining Rep sequence specificity. Schematic describing HUH-seq: an NGS-based approach for quantifying ssDNA specificity profiles of Reps. A synthetic ssDNA library containing 7 random bases (4 bases ^ 7 positions = 16,384 unique kmers) (yellow) flanked by constant regions (gray) and primer binding sites (PBS) (dark gray) are reacted with a panel of Reps, or no enzyme as a reference, in replicate, generating a two part pool containing the “uncleaved” library and the “sequence cleaved” library for each reaction. In a single PCR step, the antisense strand for the “uncleaved” pool is generated, amplified, and Nextera adapters (purple) are added with primer overhangs; the “sequence cleaved” library is not amplified due to physical separation of the PBS’s. Each set of amplicons is then barcoded with standard i7/i5 Illumina indexing sequences (green) and pooled for a single next generation sequencing run. A custom R-based analysis script generates read counts for all *k*-mers in each set of replicates, then normalizes based on total read count, and quantifies *k-*mer cleavage extent of each Rep in the panel based on fold change and percent reduction.

Moreover, the DNA in relaxase proteins is embedded in a much deeper channel than in Rep proteins. Indeed, calculations of buried solvent accessible surface area (BASA) between protein and DNA reveal a more substantial buried surface area in the binding of DNA to relaxases even when accounting for the surface area buried by the clasps (Fig 3). Both Rep and relaxase structures have obvious structurally conserved pockets in the ssDNA docking interface in which individual nucleotide bases are bound. In all structures, the sDBM, which is part of *β*1 in relaxases and *β*4 in Reps, is a major contributor to the formation of these pockets. TraI and TrwC bury nucleotides −5 and −3 in strikingly deep pockets, #1 and #2, respectively (Fig. 3). Reps have pockets in this structural region, yet they are much more shallow and only minimally bury nucleotides at −4 and −6 positions. A_-6_ is bound in the deepest of these Rep pockets, yet it is still oriented in a configuration that favors base stacking with neighboring nucleotides rather than a ‘knob-in-pocket’ interaction as seen in both TrwC and TraI structures. Conversely, both Reps have a deep pocket, #3, where the +2 cytosine base is buried, however only the TraI structure contains the positioning of the +2 base, which is interestingly not bound in this pocket (Fig. 3).

### HUH-SEQ REVEALS SUBTLE DIFFERENCES IN REP ssDNA RECOGNITION SPECIFICITY

Structural analysis of the Rep protein-DNA contact maps point to subtle differences that contribute to recognition of nearly identical nonanucleotide sequences, suggesting that Reps may differentially tolerate substitutions in the target DNA sequence. Thus, we developed a NGS-based cleavage assay approach, HUH-seq, to examine both ssDNA specificity and to explore expansion of the use of Reps in multiplexed HUH-tag applications. As a first step in assessing the ssDNA recognition specificity of Reps, we asked whether viral Rep proteins from different families and genera differentially tolerate mutations in the target nonanucleotide sequence by measuring covalent adduct formation with a standard *in vitro* SDS-PAGE cleavage assay (Table 2, Supp. Fig. 4a). However, it immediately became evident that a low-throughput assay would insufficiently characterize specificity due to widespread toleration of variable target sequences. A large number of truncations and substitutions within the nonameric sequence resulted in negligible effects on adduct formation in many cases (a full analysis of the small oligo library screen is provided, Supp. Fig. 4b and 4c). This realization prompted us to devise a high-throughput method that would reveal ssDNA recognition profiles for each Rep.

**Table 2:** Panel of 10 recombinantly expressed and purified Reps.

| HUH | Viral Species | Family | Genus | MW (kDa) | pI | cognate nonanucleotide *ori* sequence |
| --- | --- | --- | --- | --- | --- | --- |
| PCV2 | *porcine circovirus 2* | Circoviridae | Circovirus | 13.1 | 9.4 | AAGTATT*AC |
| DCV | *muscovy duck circovirus* |  |  | 12.4 | 6.1 | TATTATT*AC |
| BBTV | *banana bunch top virus* | Nanoviridae | Babuvirus | 11.2 | 9.3 |  |
| FBNYV | *faba bean necrotic yellows virus* |  | Nanovirus | 11.3 | 8.6 | TAGTATT*AC |
| WDV | *wheat dwarf virus* | Geminiviridae | Mastrevirus | 15.6 | 6.4 | TAATATT*AC |
| CpCDV | *chickpea chlorotic dwarf virus* |  |  | 14.5 | 9.1 |  |
| MSMV | *maize striate mosaic virus* |  |  | 13.4 | 7.8 |  |
| TYLCV | *tomato yellow leaf curl virus* |  | Begomovirus | 15.5 | 8.9 |  |
| CLCV | *cabbage leaf curl virus* |  |  | 13.3 | 9.1 |  |
| TGMV | *tomato golden mosaic virus* |  |  | 14.4 | 8.7 |  |

Metrics list of Rep panel used in this study describing abbreviated Rep names, viral species, family, and genus, MW and isoelectric point (pI) of the recombinant Rep domain (negating contribution from His6x-SUMO-tag), and the cognate nonanucleotide sequence from each respective viral origin of replication (*ori*).

To this end, we developed HUH-seq, an NGS-based approach used to establish comprehensive ssDNA recognition profiles of the Reps contained within a randomized ssDNA library containing 16,384 sequences, or *k*-mers. In brief, the first seven positions of the nonanucleotide target sequence are randomized in the 7N ssDNA library, where positions A_+1_ and C_+2_ are constant (“7N” - N_-7_N_-6_N_-5_N_-4_N_-3_N_-2_N_-1_*A_+1_C_+2_). The library was constrained to only seven positions in order to limit the size of the library; further design considerations are discussed in the supplementary information (Supp. Fig 5, Supp. Note 2, and Supp. Equations 1 and 2). Reps were individually reacted with the 7N ssDNA library under standard conditions and produced two populations of the library: “sequence cleaved” and “uncleaved”. A primer set containing Nextera adapters was used to generate the antisense strand and to amplify the “uncleaved” population in a single PCR step, while the “sequence cleaved” population remained unamplified. The “uncleaved” amplicons were barcoded with standard dual-indices and sequenced using the HiSeq platform to obtain read counts for every sequence in the library (*k*-mer). Read counts from reference replicates (no Rep added to the reaction) were used to calculate log_2_-fold-change (FC) and read count percent reduction based on the difference between the normalized reference library read counts and normalized “uncleaved” read counts for each Rep treatment (Fig. 4).

We generated weighted sequence logos based on a *k*-mer reduction analysis with a threshold value of 0.3 or greater to reduce noise (Fig. 5). Percent reduction for each *k*-mer was calculated by comparing the normalized *k*-mer read counts for each Rep treatment in triplicate to *k*-mer read counts from the reference library. For each position in a Rep sequence logo, individual characters were scaled by the average percent reduction of all *k*-mers containing that character and position. Because every sequence permutation 5’ of the *nic* site is present in the 7N ssDNA library, sequence logos reveal Rep preferences for nucleotides relative to one another. The most obvious result is that the most preferred nucleotides in the first seven positions of sequence logos are nearly identical to the cognate nonanucleotide *ori* sequence found in each respective viral genome (Fig. 5). While it is not surprising that the preferred target sequence is the same as the cognate nonanucleotide *ori* sequence cleaved *in vivo*, it also gives high confidence that HUH-seq can be used to quantitatively rank the *k*-mers cleaved by each Rep, analyze patterns that dictate these ssDNA recognition profiles, and further characterize differences between individual Reps.

**Fig. 5:**
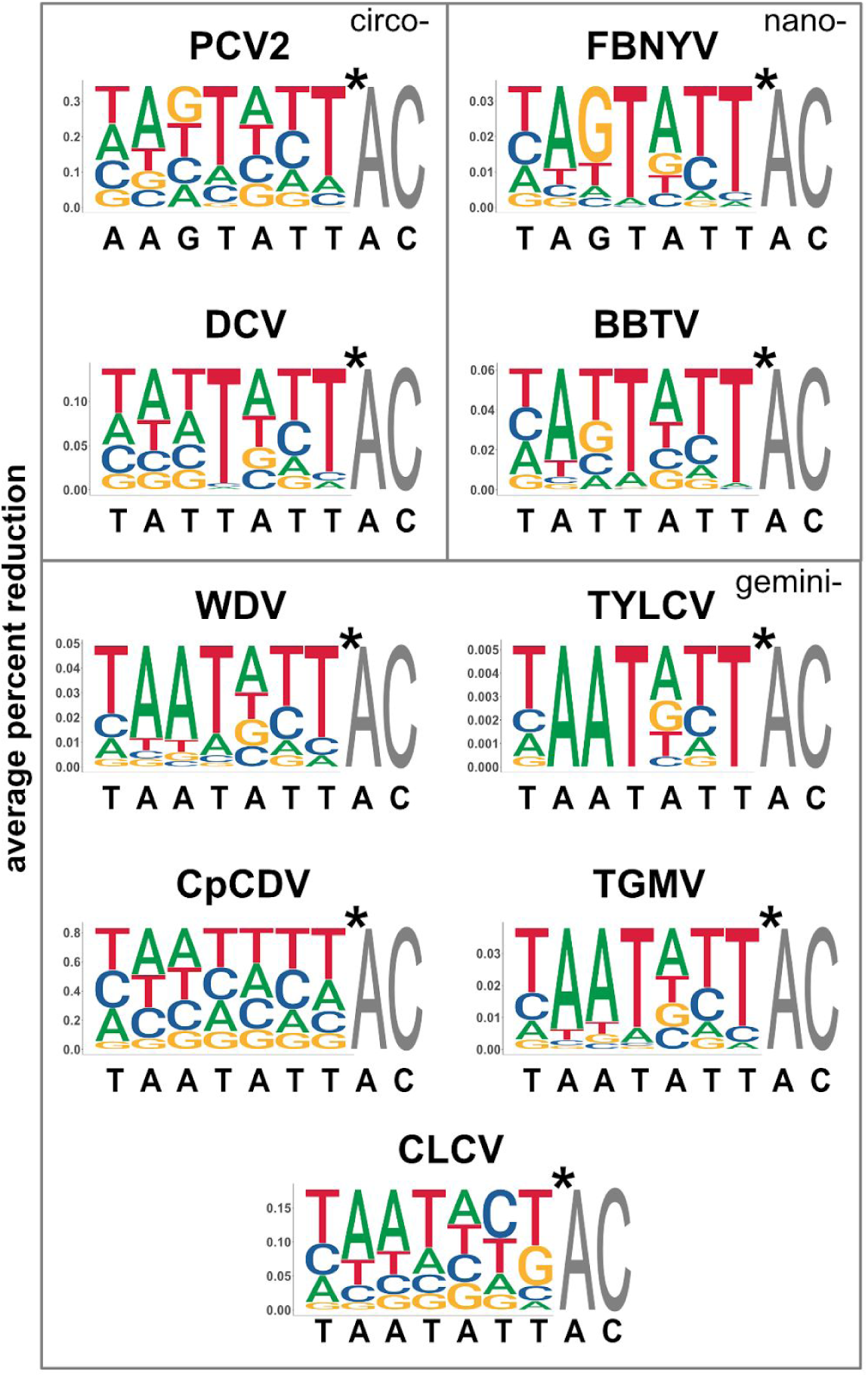
Weighted sequence logos generated from HUH-seq cleavage data. Weighted sequence logos for nine of the ten Reps based on percent reduction generated using ggseqlogo with values under 0.3 set to 0.0 in order to remove noise obtained from the HUH NGS cleavage assay. Heights are scaled to represent the average percent reduction of each base at each position when compared to the reference library. Sequences in black below each logo are the cognate nonanucleotide *ori* sequences from each respective virus. Logos are organized by viral families as labeled inside the gray boxes.

Within each sequence logo, there are differentially preferred nucleotide positions. Positions T_-4_ and T_-1_ are almost unanimously the most preferred, while conservation of the A_-3_ and T_-2_ positions has a relatively low impact on cleavage. There are also discernible trends between Reps from different families. For example, geminivirus Reps have a strong preference for adenine at the −5 position, whereas Reps from other families prefer thymine or guanine there (Fig. 5). The y-axis scale of the weighted sequence logos also indicates the relative overall cleavage efficiency between Reps. For instance, PCV2 has a maximum average percent reduction of about 0.35 and cleaves roughly 10-fold more sequences than FBNYV, which has a maximum value of about 0.035. This indicates that PCV2 ssDNA recognition is more promiscuous than that of FBNYV. CpCDV has the highest maximum average percent reduction of 0.8 and has minimal nucleotide preference, indicating that it has the most relaxed sequence specificity (Fig. 5). Other considerations and caveats of HUH-seq analysis are discussed in supplemental information (Supp. Note 4 and Supp. Fig. 6).

### REP ssDNA RECOGNITION PROFILES CORROBORATE STRUCTURAL OBSERVATIONS

Next, we quantified and assigned contributions of the ssDNA docking interface in the Rep structures to each nucleotide using DNAproDB by calculating the BASA as well as the total number of protein-DNA contacts (the sum of hydrogen bonds and Van der Waals interactions within 4 Å). Figures 6a and 6b summarize the total BASA for and the total number of contacts with nucleotides corresponding to the cognate nonanucleotide *ori* sequence either with the entire nucleotide or the base only. These measurements in combination with the ssDNA recognition profiles of WDV and PCV2 were used to search for structural reasons why nucleotides in certain positions of the target sequence are conserved. A comprehensive table containing BASA and contact values of each of the three structures featured in this study is also provided (Supp. Table 1). As expected, higher BASA values generally correlated to high numbers of contacts.

**Fig. 6:**
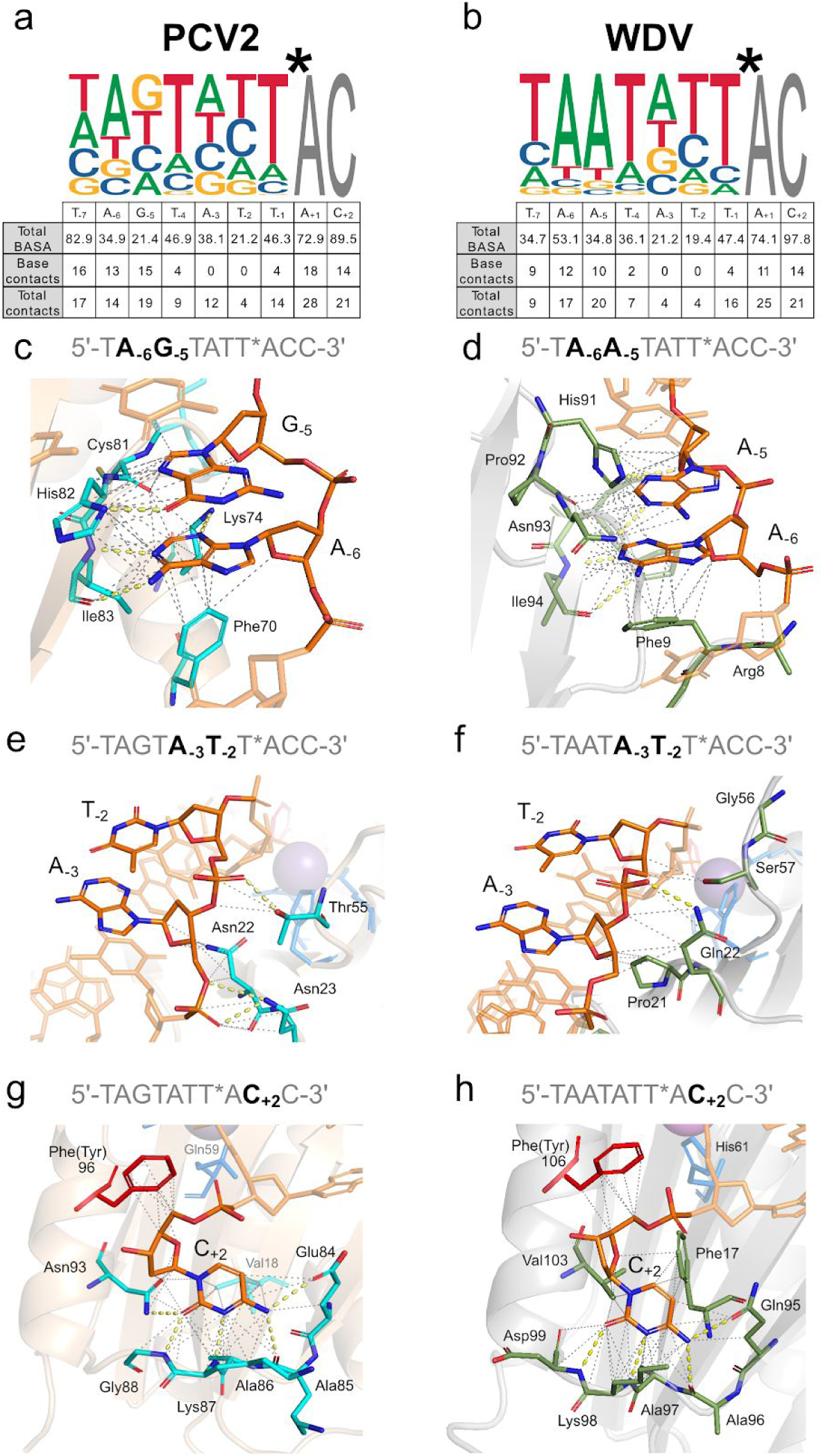
Comparison of Rep protein-DNA interactions and HUH-seq specificity profiles. **a**,. PCV2 + 10mer (6WDZ) and **b**,. WDV + 10mer (6WE0) BASA values and total number of protein-DNA contacts compared to weighted sequence logos from HUH-seq analysis. Both polar and van der Waals interactions are counted within 4 Å. **c-h**, Atomic interactions between highlighted nucleotides within the bound 10-mers of PCV2 and WDV structures are shown with yellow dashes for polar contacts and gray dashes for van der Waals interactions within 4 Å. PCV2 cartoon is depicted in beige and residues interacting with DNA as sticks shown in cyan colored by atom. WDV cartoon is depicted in gray and residues interacting with DNA as sticks shown in green colored by atom. The PCV2 Phe96 and WDV Phe106 represent the catalytic tyrosine as sticks in red, divalent ion coordinating residues are in blue, and the manganese ion is a sphere in magenta. The 10-mer is shown as sticks in orange and colored by atom as highlighted in the panel.

PCV2 bound 10-mer and WDV bound 10-mer structures have a similar total number of residues contacting DNA, 28 and 26 residues, respectively and have a high concentration of base contacts and total contacts near the 5’ and 3’ termini of the 10-mers (Fig. 6a and 6b). In Figure 6c - 6h, significant structural differences are highlighted between the contacts of nucleotides at different positions for both PCV2 (c, e, and g) and WDV (d, f, and h). A_-3_ and T_-2_ are the least conserved nucleotides at their indicated position. This is structurally consistent because there are zero contacts with the bases for both PCV2 and WDV indicating that specific nucleotides are not as prefered at these two positions because the interactions are exclusively with the ribose and phosphate of the nucleotides (Fig. 6e and 6f). The 10-mer bound to PCV2 differs at position −5 between guanine and adenine with respect to the 10-mer bound to WDV. His91 and Asn93 of WDV facilitate polar contacts with A_-5_, which may give WDV more specificity at position −5, whereas there is only one polar contact with G_-5_ by His82 in the PCV2 structure, which results in less stringent specificity. (Fig. 6c and 6d). Finally, in both structures C_+2_ dwells in a pocket of the protein surface with the highest BASA and total contact values (Fig. 6g and 6h). Eight residues have contacts with C_+2_ in both structures, and five of these residues make up the last positions of the sDBM.

In contrast, T_-4_ is highly conserved as evident in all Rep ssDNA recognition profiles, but we observed only a marginal number of protein contacts with the base itself (Fig 6a and 6b). We hypothesize that the WC base pairing of T_-4_ with A_+1_ is a major contributor to the U-shaped conformation rather than contributing to sequence specificity via residue interactions with the base. Though Reps exhibit interactions with bases that contribute to specificity, it is clear from the ssDNA recognition profiles and minimal protein-DNA contacts at certain positions that Rep cleavage is also promiscuous, cleaving a wide range of target sequences.

### DISCOVERING INTRINSIC ORTHOGONAL REP TARGET SEQUENCES USING HUH-SEQ

During initial assessments of the HUH-seq analysis results, we noticed that there were individual target *k*-mers with drastically different log_2_FC values between different Rep protein treatments. This prompted us to ask whether we could identify pairs of *k*-mers that would allow us to selectively label two Reps in a single reaction mixture with unique oligos. For instance, *k*-mer, AGTCAAT (#2884) has a log_2_FC value of −3.44 for PCV2 and a near zero log_2_FC for every other Rep (Supp. Fig. 7). This result was validated using the standard in vitro cleavage assay by reacting PCV2 Rep with a synthetic oligo containing this *k*-mer sequence. Indeed, only PCV2 formed a covalent adduct with the oligo harboring this target sequence (Supp. Fig. 7). Interestingly, this target sequence contains 4 substitutions with respect to the circovirus *ori* sequence at positions −6, −5, −4, and −2, again highlighting the promiscuous nature of Reps. This result revealed that searching for combinations of Reps and *k*-mers may result in the discovery of naturally occurring orthogonality despite cross-reactivity between cognate nonanucleotide *ori* sequences.

To explore the possibility of naturally occurring orthogonality between two Reps, we wrote a script to extract pairs of *k*-mer sequences and Reps predicted to lack cross-reactivity based on log_2_FC values. Figure 7a displays a summary heatmap of the number of such *k*-mer pairs existing for every set of Rep pairs, based on threshold values of −0.3 log2 FC and greater (likely forming no adduct) and −3.0 log_2_FC and lower (likely having high adduct formation). In one example, we identified the *k*-mer sequence, CATTTCT (#5112), in which DCV had a −4.13 log_2_FC and WDV had a −0.33 log_2_FC, and another *k*-mer sequence, TAAATCT (#12344), in which DCV had a −0.20 log_2_FC and WDV had a −4.11 log_2_FC, indicating orthogonality between DCV and WDV for these two *k*-mers. We validated this observation with a standard in vitro cleavage assay including a short time course with 1, 5, and 10 minute time points. DCV formed about 97% adduct with a synthetic oligo harboring *k*-mer #5112 over the course of 5 min, and WDV formed about 62% adduct with a synthetic oligo harboring *k*-mer #12344 over the course of 10 min. As expected, no cross-reactivity was observed between WDV with *k*-mer #5112 or DCV with *k*-mer #12344 (Fig. 7a).

**Fig. 7:**
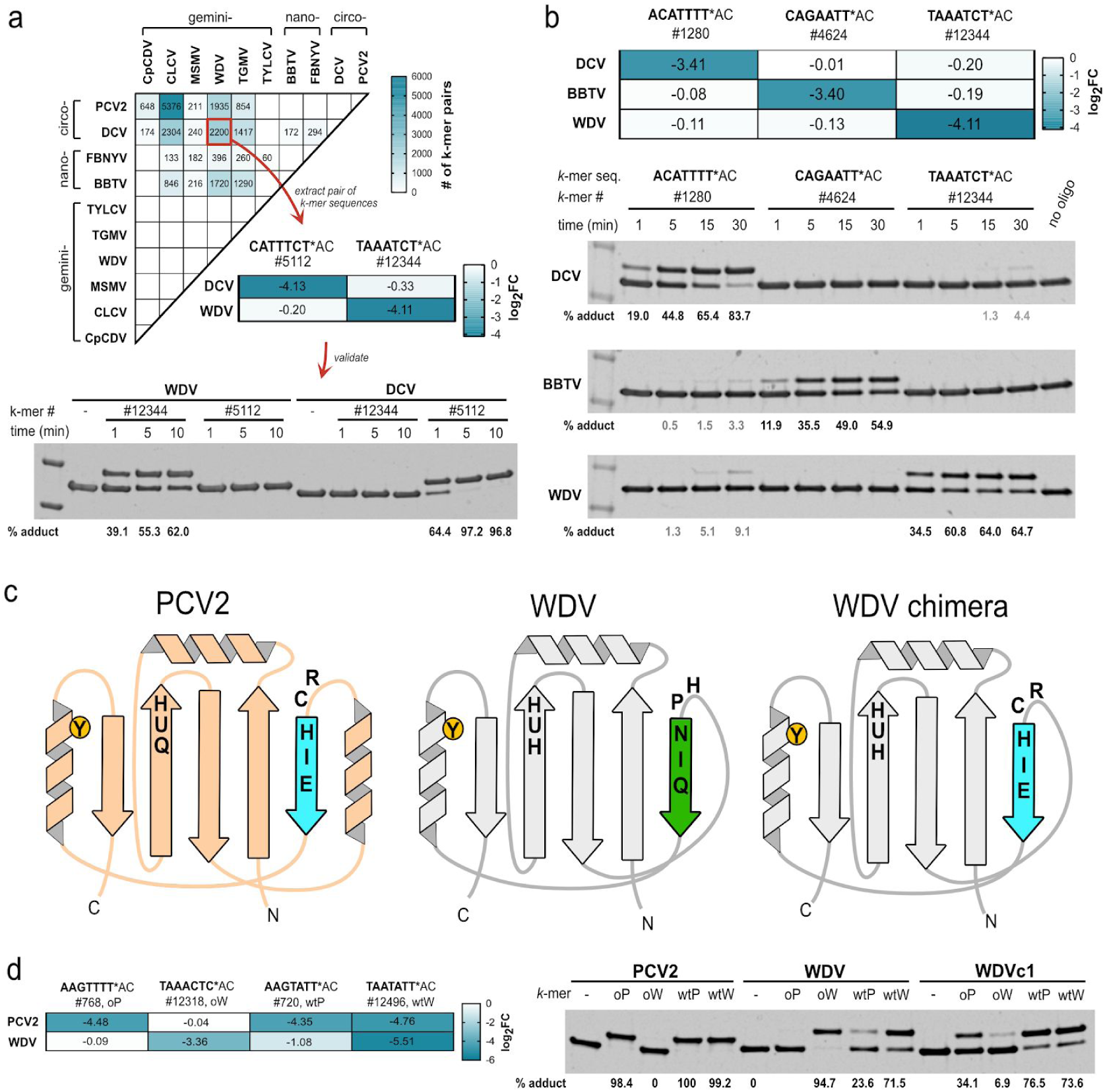
Discovery of orthogonal Rep target sequences and rational engineering of Rep specificity. **a**, Heatmap displaying the number of *k*-mer pairs for a specific Reps set likely to be orthogonal using an asymmetric log2FC threshold based on values from HUH-seq analysis, blank cells indicate zero such *k*-mer pairs. The threshold values are set to log2FC values greater than −0.3 (indicating no *k*-mer cleavage) and log2FC values less than −3 (indicating high *k*-mer cleavage). Each cell of the heatmap represents the total number of possible *k*-mer pair combinations likely to be orthogonal for a particular set of two Reps. Sets are based on this asymmetric threshold in which the first Rep in the set has high cleavage of one *k*-mer in the pair and no cleavage of the other *k*-mer in the pair - vise versa for for the second Rep in the set, indicating orthogonality. As an example, one *k*-mer pair (#5112 and #12344) of the 2200 possible combinations indicated by the WDV vs. DCV cells were synthesized in the context of the flanking regions of the 7N ssDNA library, and cleavage orthogonality was validated using the standard HUH in vitro cleavage assay. Recombinant WDV and DCV were reacted under standard conditions over a short time course with synthetic oligos harboring *k*-mers sequences #5112 or #12344. Percent covalent adduct was calculated. **b**, Set of three Reps and corresponding orthogonal set of three *k*-mer sequences as indicated by log2FC values from HUH-seq analysis. Oligos synthesized harboring *k*-mer sequences (#1280, #4624, #12344) were reacted with DCV, BBTV, and WDV recombinant Reps at room temperature with 1.5x molar excess oligo to Rep protein over a short time course. **c**, A schematic illustrating the construction of the WDV chimera (WDVc1) containing the first five amino acids of the PCV2 sDBM. **d**, The heatmap displays HUH-seq log2FC values for PCV2 and WDV reactivity with cognate nonanucleotide sequences (*k*-mers #720 and #12496) and a pair of *k*-mers (#768 and #12318) predicted to react orthogonally. The *k*-mers #720 (wtP), #12496 (wtW), (#768 (oP), and #12318 (oW) were synthesized in the context of the flanking sequences of the 7N ssDNA library and reacted in a 5x molar excess with PCV2, WDV, and WDV chimera (WDc1) recombinant protein for 30 min and 37°C.

We next searched for triple orthogonal sets of Reps from our panel. As an example, the set containing *k*-mer sequences #1280, #4624, and #12344 are predicted to react orthogonally with DCV, BBTV, and WDV recombinant Reps, respectively, as indicated by log_2_FC values (Fig. 7b). Similar to our method for validating double orthogonal sets, we tested the orthogonality of this set using the standard in vitro cleavage assay and calculated percent adduct formed with each combination of *k*-mer and Reps over a short time course. Expected orthogonality was achieved with over 50% covalent adduct formation after 30 minutes for each of the three Reps with 0 - 9% crossreactivty identified (Fig. 7b).

Notably, 23 of the 28 Rep sets from different viral families contained significant *k*-mer pairs likely to be orthogonal, yet there were no instances of orthogonal *k*-mer pairs for Reps derived from the same viral family (Fig. 7a). Hence, the ssDNA binding moieties of Reps within the same family may be too similar to yield orthogonal adduct formation. This is a curious result in the case of DCV and BBTV where 294 potentially orthogonal *k*-mer pairs were identified that are from different Rep families but which recognize identical cognate nonanucleotide *ori* sequences (Fig. 7a). This indicates that perhaps DCV and BBTV use different interactions to recognize the same cognate sequence allowing for divergent specificity at each nucleotide position. Indeed, 6 out of 9 residues in the sDBM are different between DCV and BBTV Reps (Fig. 4c). Using HUH-seq, we can pick out subtle differences in Rep specificity in order to extract double and triple orthogonal *k*-mers and Reps sets that can be used in multiplexed HUH-tag technologies. This could potentially negate the necessity for or the creation of ways to use Reps in combination with the larger and slower relaxases or commercial fusion tags.

### RATIONAL DESIGN OF A WDV CHIMERA CONFERS PCV2-LIKE SEQUENCE SPECIFICITY

The identification of the sDBM, which we hypothesized was responsible for sequence specificity in Reps as well as the discovery of pairs of target sequences between two Reps that should not cross-react, inspired us to swap sDBM residues between two Reps and ask if we could alter sequence specificity. We swapped out the first five amino acids of the WDV sDBM for those of PCV2, creating a WDV chimera (WDVc1) as a proof-of-concept that Rep specificity could be altered by rational design in a predictable manner (Fig. 7c). Because many of the amino acid side chains in both Rep structures have direct contacts with bases at the 5’ end of the ssDNA, we hypothesized WDVc1 would have sequence specificity more closely reflecting that of PCV2. First, we identified a pair of predicted target sequences for PCV2 and WDV using HUH-seq, where WDVc1 reacts with the *k*-mer #768 (oP), which was predicted to only react with PCV2, to a greater extent than *k*-mer #12318 (oW), which was predicted to only react with WDV (Fig. 7d). Similar to PCV2, WDVc1 reacts robustly with the cognate nonameric sequence of PCV2, *k*-mer #720 (wtP), as well as with the cognate nonameric sequence of WDV, *k*-mer #12496 (wtW), (Fig. 7c). Thus, we show how the sDBM is a key feature of Reps that may be rationally engineered to predictably alter sequence specificity.

## Discussion

Our previous work using Reps from circular Rep-encoding ssDNA (CRESS-DNA) ^41,42^ viruses in HUH-tagging illustrated that while these small proteins are extremely efficient at forming covalent adducts with ssDNA compared to their relaxase counterparts, Reps from related circular ssDNA viral families exhibit cross-reactivity due to a combination highly similar DNA target sequences and promiscuous specificity ^4^. However, the lack of molecular details about how DNA recognition is encoded in viral Reps has prevented exploration of their potential for use in multiplexing applications or engineering for sequence specificity. Elucidating the molecular underpinnings of DNA processing by Reps could lead to new antiviral therapies for preventing Rep-mediated viral infections in plants and humans, enhance HUH-tag multiplexing for emerging DNA-labeling applications, and allow engineering of designer sequence specificity for new gene therapy tools.

We solved the first co-crystal structures of Rep HUH-endonucleases in complex with ssDNA target sequences and developed a complementary NGS cleavage assay, termed HUH-seq, which allowed us to: 1) identify a key structural motif called sDBM that largely encodes sequence specificity between Rep families; 2) harness subtle differences in DNA recognition to discover target sequences that selectively react with a Rep from one family and not with Reps from other families; and 3) perform proof-of-concept engineering of a few amino acids to switch DNA recognition specificity of WDV to that of resemble PCV2.

We first determined the molecular basis of DNA recognition of viral Reps by solving crystal structures of viral Reps with ssDNA containing the cognate *ori* sequence trapped in the pre-cleavage conformation. Several apo structures of viral Reps in the absence of DNA have been reported ^39,40,43–45^ as well as parvovirus AAV5 Rep structures bound to distal auxiliary regions of dsDNA with the inverted terminal repeat involved in Rolling Hairpin Replication ^46,47^. However, the Rep structures presented here for the first time illuminate the interface responsible for specific ssDNA recognition necessary for ssDNA processing. The most striking feature of the ssDNA bound structures is the central role played by a motif we call the sDBM. This motif is highly conserved between members of the same viral Rep family but divergent between families, yet it maintains its key function of binding ssDNA for cleavage across the entire HUH endonuclease superfamily. (Fig. 3, Supp. Fig. 3). The sDBM partially overlaps with a previously identified ∼20 amino acid long motif conserved in the geminivirus family called the GRS, which was hypothesized to interact with DNA ^48^. The sDBM has two critical roles: first, it promotes a general task of priming the conformation of the ssDNA substrate in a pre-cleavage U-shape, and second, it provides specific protein-ssDNA contacts, using both peptide backbone and sidechain contacts, to confer specificity for ssDNA based on amino acid sequence. The structures reveal a divergent network of protein-DNA contacts maintained by the sDBM clustered around the −5 position of the ssDNA that contains the substitution between nearly identical target sequences.

Looking more broadly at the Rep-DNA interactions in the context of the entire HUH-endonuclease superfamily, the sDBM is apparently a ubiquitous motif contributing to DNA recognition. This is illustrated by available relaxase structures captured in the pre-cleavage state ^17,28–30^. In Reps, the sDBM is located in the middle of the structure and consists of the fourth beta strand and a portion of the preceding loop. In relaxases, however, the sDBM is located at the extreme N-terminus due to the circular permutation of relaxases with respect to viral Reps. Thus amino acids derived from a nearby C-terminal loop likely contribute to sDBM’s role in relaxases ^49^. The sDBM in relaxases also plays a major role in forming specific contacts especially with nucleotides bound in deep pockets of the protein surface. The fact that the N-terminus plays such a critical role in binding DNA has a negative impact on the use of relaxases as C-terminal fusion tags. Indeed, some relaxases cannot bind to DNA unless the N-terminus is free ^4^. Moreover, the “knob-in-hole” mode of DNA recognition in which certain DNA bases are buried in deep pockets on the surface of the proteins, while conferring higher specificity ^32^ than the shallow pockets found on Reps, makes engineering of the protein for recognition of different bases all the more daunting. In contrast to Reps, relaxases also have additional “clasp” and DNA hairpin recognition motifs that hold the DNA in place that contribute to the higher specificity relaxases have for their target sequences. In nature, there are several reasons why Rep specificity could be more promiscuous than that of relaxases. If conjugative plasmid transfer occurs at an erroneous origin, it could result in catastrophic consequence for the recipient organism’s fitness; whereas, pressure for a virus to initiate replication of a small genome or to integrate inside a host’s genome at a very specific sequence is minimal ^32^. In addition, relaxases may have higher specificity for the DNA sequence 5’ of the *nic* site for more efficient catalysis of rejoining of the free 3’OH of the DNA post-transfer, whereas RCR resolution would likely require a second dimerizing Rep for termination ^50^. It should also be noted that Rep specificity could also simply be constrained due to a smaller interface surface area due to limited gene size, as most viral Rep genomes are under five kilobases ^41^.

Characterization of the Rep/ssDNA interface provides a platform to model other Rep ssDNA interfaces as well as an avenue to explore disruption of the interface for antiviral treatments of plant and human Rep-mediated viral infections ^51^. Billions of dollars worldwide are lost in agriculture every year from the decimation of crops such as tomatoes, cassava, cotton, and beans by geminivirus infection ^52^. In a human disease context, parvovirus B19 human infections can lead to serious or fatal outcomes for a fetus ^53,54^ and are associated with autoimmune diseases in adults ^21,22^. This has sparked a push to develop treatments and vaccines ^55^. Present antiviral strategies, both viral protein interfering and gene silencing approaches, are either only minimally effective or are eventually subverted by conferred resistance from a rapidly evolving viral genome ^51,56–60^. However, development of antivirals specifically targeting the ssDNA binding of Reps could more effectively retain long-term resistance.

The Rep structures revealed highly conserved protein-DNA interfaces with subtle differences that prompted us to ask whether Reps within families and from different families differentially tolerate mutations of the cognate *ori* sequence. One reported a relatively high throughput strategy for querying key nucleotides in bacterial conjugation mediated by the relaxase TrwC that used saturation mutagenesis in concert with a functional DNA-transfer readout ^32^, which is not a feasible readout for Reps. Other NGS-based ssDNA recognition approaches, for example of cytosine deaminases ^61^ or DNA aptamer-binding protein targets ^62^, also use direct sequencing readout methodology. However, a direct readout of Rep cleavage is technically challenging due to the need to amplify physically separated cleaved DNA molecules and covalent attachment of the Rep to the new 5’ end of the cleaved molecules. Instead, HUH-seq, allows for the quantitative readout of Rep cleavage specificity using a ssDNA library with a subtractive, or reduction, readout.

The most obvious result of the assay and validation of the HUH-seq approach was the appearance of the known cognate *ori* sequence as the most preferred cleaved sequence for most Reps tested. Moreover, many aspects of HUH-seq analysis corroborate the structural analysis, notably that positions −5 and −6 confer the most specificity via protein residue and nucleotide base contacts, while nucleotide positions −2 and −3 conferred the least specificity due to a lack of contacts with nucleotide base contacts. Excitingly, we found HUH-seq can be used to distinguish subtle differences between Rep nucleotide preferences despite overall lack of specificity, so much so that intrinsic orthogonality between non-cognate target sequences can be extrapolated between Reps from different, yet closely related, viral families with highly similar or even the same cognate nonanucleotide *ori* sequences.

The finding of oligos with intrinsic orthogonality between Rep families demonstrates the feasibility of using Reps in multiplexing applications without the need for protein engineering, despite their apparent promiscuity. Moreover, there are currently 10 other CRESS-DNA families yet to be explored with HUH-seq ^41^, meaning that multiplexing could be expanded to up to 13 Reps in a given system (i.e. have 13 HUH-fusions in an application such as DNA barcoding of proteins of interest and add 13 DNA barcodes that should specifically react with only one given Rep HUH-tag). DNA-tagging is the basis of established technologies such as proximity ligation ^63^ and DNA-PAINT super-resolution imaging ^64^ as well as emerging applications such as multiplexed single-cell proteomics ^65^, and optics free DNA microscopy ^66^, where parallel tracking of proteins occurs using NGS. It is of note that HUH-fusions would allow conjugation of oligos to ScFv’s and increasing popular nanobodies, which could expand the utility of many of these applications which utilize oligo-conjugated antibodies. Because HUH-tag linkages are specific and covalent, can occur intra- or extracellularly without additional reagents, and are now multiplexable, they are ideal fusion tags for these applications.

With minor alterations, we foresee the utility of the simple HUH-seq approach for the sensitive detection of sequence specificity profiles for enzymes such as dsDNA nucleases by simply using a dsDNA library, RNA-cleaving enzymes by adding a single reverse transcriptase step, or site-specific nucleotide modifying enzymes by relying on a covalent modification that blocks PCR amplification. The existing high-diversity library methods used to determine the dsDNA specificity of zinc finger nucleases ^67^, Cas9 ^68^, transcription activator-like effector nucleases (TALENs) ^69^, and other restriction enzymes ^70^ are powerful and direct cleavage readout approaches, However they require a number of extra library preparation steps and may be limited to only dsDNA libraries. Additionally, if sequence binding, rather than cleavage, could be optimized as a readout, HUH-seq could be developed as a facile alternative method to approaches such as SELEX-seq ^71^ and could determine binding sequence preference of shorter DNA binding motifs of enzymes such as transcription factors, eliminating the need for multiple rounds of sequence enrichment.

While subtle but specific family differences in DNA recognition coupled with HUH-seq permits modest viral Rep multiplexing, expanded multiplexing capability could be achieved by engineering the protein to recognize designer DNA sequences. Engineering one HUH-endonuclease to react with another HUH-endonuclease target sequence has been demonstrated for AAV, but it involved swapping large protein domains ^72^. Similarly, a double mutant of the TraI relaxase that conjugates the F plasmid was able to switch specificity to the related R100 plasmid target sequence, though the engineering was performed by testing and mutating all distinct amino acids residues between the two relaxases ^49^. We have shown in an elegant example of rational engineering that by simply mutating four amino acids within the sDBM of WDV and PCV2, specificity can be predictably altered. This approach was made possible not only by structural insights, but also because we can identify intrinsically orthogonal target sequences that would react specifically with each Rep using HUH-seq.

Predictable altering of ssDNA specificity by targeting the residue composition of the sDBM either by rational design or directed evolution could motivate development of engineered HUH-tags with defined sequence specificity to facilitate massively parallel Rep-based applications. Further, HUH endonuclease mediated genome integration is severely constrained to a small number of sites. For example, AAV Reps and relaxases could be engineered to integrate into a broad range of desirable sites to greatly expand the usefulness of this critical gene therapy tool. HUH-seq will be an indispensable approach in defining the specific profiles these engineered HUH-endonucleases have for their designer target sequences.

There are several potential improvements on our studies. While these Rep-ssDNA structures are likely highly representative of RCR mediating ssDNA viruses, parvovirus Reps exhibit some sequence and structural differences. For example, the sDBM of AAV Reps include an additional charged loop and have been shown to also contact the ITRs perhaps giving it an added level of specificity ^47^. Elucidating the exact ssDNA binding mode of action of parvovirus Reps would provide more specific information for engineering specificity for gene integration applications or designing Rep-targeting antivirals for human disease causing viruses. HUH-seq is limited by the diversity size of the library; however, as NGS read capacity, speed, and cost-effectiveness increase, along with computational processing and data storage, library size may become a negligible shortcoming of HUH-seq. While we use a limited diversity library of 7 randomized nucleotides in this study, it still allowed us to interrogate specificity of many Reps at once in addition to allowing a number of controls in a single HiSeq sequencing lane. A more complete Rep specificity profile using an expanded sequence library is possible using HUH-seq, however this greatly limits the number of Reps and controls that can be used in a single NGS sequencing lane. Using multiple sequencing runs and lanes would be a simple solution to this issue though would dramatically increase cost.

Together the combination of structural and NGS approaches demonstrate that viral Reps, with desirable size and reaction efficiency but low apparent sequence specificity, can be exploited in multiplexing applications by engineering DNA target sequences and protein sequences. These findings will drive further studies into engineering HUH-endonuclease recognition of ssDNA and expanded application of HUH-endonucleases as HUH-tags.

## Methods

### MOLECULAR CLONING, PROTEIN EXPRESSION, AND PURIFICATION

The N-terminal domain of all Reps were synthesized as E. coli codon-optimized gene blocks from Integrated DNA technologies (IDT) and designed with 15 nucleotides on each end that were homologous to regions of the linearized pTD68/His6-SUMO parent vector digested with BamHI and XhoI. Final His6-SUMO-Rep constructs were created with the In-Fusion HD Cloning Kit (Takara) and sequence confirmed with Sanger sequencing (Genewiz). Purified plasmids were transformed into BL21(DE3) E. coli competent cells (Agilent), initially cultured in 1L LB broth at 37°C, then induced at OD600 with 0.5 mM IPTG (isopropyl-d-1-thio-galactopyranoside, Sigma Aldrich), and then grown for 16 hours at 18°C. Collected cell pellets were resuspended in 10 mL of lysis buffer (50 mMTris pH 7.5, 250 mM NaCl, 1 mM EDTA, complete protease inhibitor tablet (Pierce)) and pulse sonicated for several one minute rounds. The suspension was centrifuged at 24,000xG for 25 min, and supernatants were batch bound for 1 hour with 2 mL HisPure Ni-NTA agarose beads (ThermoFisher) and equilibrated with wash buffer (50 mM Tris pH 7.5, 250 mM NaCl, 1 mM EDTA, 30 mM imidazole). After lystate cleared the gravity column, beads were washed with 30 mL wash buffer, and proteins were eluted from gravity columns with elution buffer (50 mM Tris pH 7.5, 150 mM NaCl, 1 mM EDTA, 250 mM imidazole). Protein was further purified and buffer exchanged into 50 mM Tris pH 7.5, 150 mM NaCl, 1 mM EDTA using the ENrich SEC70 (Bio-Rad) size exclusion column. Aliquots were stored at −20°C and −80°C at 30 μM. SUMO-cleaved recombinant PCV2^Y96F^ and WDV^Y106F^ stocks for crystal screening were prepared in a similar manner as above, however Ni-NTA fractions were dialyzed into 50 mM Tris pH 7.5, 300 mM NaCl, 1 mM EDTA as above with the addition of 1 mM DTT and SUMO-cleaving protease ULP-1 at 5 U per 1 L of E. coli overnight at 4°C. Dialyzed samples were batch bound a second time with Ni-NTA beads and were flowed through a gravity column to remove cleaved His6-SUMO and His6-ULP-1. Protein was concentrated with spin concentrators (Amicon Ultra-15 Centrifugal Filter Unit, 3 kDa cut-off) to 16 mg/mL.

### CRYSTALLIZATION, DATA COLLECTION, AND PROCESSING

An 8-mer oligonucleotide (5’-dAATATTAC-3’) from part of the geminivirus origin of replication sequence was reconstituted in ddH20 at 10mM and mixed with recombinant WDV^Y106F^. We used Rigaku’s CrystalMation system to perform a broad, oil-immersion, sitting drop screen of the protein-DNA mixture in the presence of either magnesium or manganese. Crystals were achieved using 8 mg/mL protein solution containing 1.1-fold 8-mer and 5mM MnCl2 with a well solution of 12% (w/v) PEG8000 precipitating agent, 0.2 mM zinc acetate, and 0.1 M sodium cacodylate at pH 6.5. The crystals belong to space group *P* 4_1_ with unit cell dimensions of *a* = *b* = 50.63 Å, *c* = 241.98 Å. Four complexes were present per asymmetric unit. *P* 4_1_ 2_1_ 2 was also a solution for this crystal morphology with 2 complexes per asymmetric unit, however, we failed to solve a suitable model at higher symmetry possibly due to variations in the crystal structure. Addition of any cryoprotectant to these crystals resulted in poor diffraction; the crystals seemed to collapse upon vitrification. Our solution to this issue was to collect datasets using an in-house, x-ray diffractometer (Rigaku Micromax-007 Rotating Anode, Rigaku Saturn 944 CCD Detector) at room temperature. Radiation caused minimal crystal damage, and over 100 frames could be obtained from a single crystal. All data was processed with the HKL suite.

WDV^Y106F^ + 10-mer crystals were also obtained with 1:1 protein solution to well solution, where the well solution was constant (12% (w/v) PEG8000 precipitating agent, 0.2 mM zinc acetate, and 0.1 M sodium cacodylate at pH 6.5), and contained 1mM 10-mer oligonucleotide (5’-dTAATATTACC-3’). Protein and MnCl2 concentration, 8 mg/ml and 5 mM respectively, were also held constant. Crystals were soaked in 25% glycerol, and a dataset was collected at the APS Beamline 24 (NE-CAT). Crystals diffracted to 1.8 Å and belong to the *P* 2_1_ 2_1_ 2_1_ space group with unit-cell parameters: *a* = 45.57 Å, *b* = 50.01 Å, *c* = 73.44 Å. One complex was present per asymmetric unit.

We also used Rigaku’s CrystalMation system’s broad, sitting drop screen to identify potential conditions for PCV^Y96F^ + 10-mer crystallization. The protein solution contained 8 mg/ml protein, 1 mM 10-mer oligonucleotide (5’-dTAGTATTACC-3’), and 5 mM MnCl2. Small needle crystals were obtained with 1:1 protein solution in a well solution of 0.1 M ammonium acetate; 25% polyethylene glycol 3,350; 0.1 M Bis-Tris pH 7. Crystals were soaked in 25% glycerol, and a dataset was collected at the APS Beamline 24 (NE-CAT). Crystals diffracted to 1.93 Å and belong to the *P* 64 space group with unit-cell parameters: *a* = *b* = 99.61 Å, *c* = 73.72 Å. There were 3 complexes per asymmetric unit.

### STRUCTURE SOLUTION AND REFINEMENT

The WDV^Y106F^ + 8-mer structure was solved with the molecular replacement function in PHENIX using our previously solved structure of apo WDV Rep (PDB ID: 6Q1M) as a model. We visualized the electron density map using Coot ^73^ and recognized clear patches in the map to be a chain of nucleotides. All 8 nucleotides of the 8-mer oligonucleotide were unambiguously built into well-defined electron density of each of the 4 complexes in the asymmetric unit. Subsequent refinement was performed with default settings of PHENIX auto.refine with NCS applied ^74^ and alternated with visual inspection and model correction. Final R-work and R-free were 0.167 and 0.236 respectively.

The WDV^Y106F^ + 10-mer, *P* 2_1_ 2_1_ 2_1_ structure was solved with the Phaser molecular replacement function in PHENIX using the previously solved WDV^Y106F^ + 8-mer structure. The two additional nucleotides were modeled into appropriate density. Again, Coot was used for model building, and PHENIX auto.refine was used for refinement. The final R-work and R-free were 0.173 and 0.224 respectively.

A model for molecular replacement was generated in PyMol by superimposing the WDV^Y106F^ + 8-mer structure with the porcine circovirus Rep domain (PDB ID: 5XOR) structure. The 8-mer from the WDV model was added to the PCV Rep domain model and used for Phaser molecular replacement in Phenix. The two additional nucleotides were modeled, and the oligonucleotide sequence was corrected using Coot. PHENIX auto.refine was used for refinement. While refining, two of the complexes in the asymmetric unit had well-defined electron density. Density corresponding to the third complex was poorly defined, and modeling was difficult. As a result, R-values are higher than normal for this resolution structure. R-work and R-free were calculated to 0.245 and 0.301 respectively.

### STANDARD IN VITRO HUH CLEAVAGE ASSAY

Cleavage of the synthetic oligos was carried out using final concentrations of 3 µM SUMO-Rep and between 4.5 - 30 µM oligo in 50 mM HEPES, 50 mM NaCl, and 1 mM MnCl2 for 30 min at 37°C. The reactions were quenched with 4x Laemmli buffer containing 5% β-ME, boiled for 5 min at 100°C, and run on a 4-12% SDS-PAGE acrylamide gel. For time course reactions, aliquots were removed from an HUH reaction master mix at specified time intervals and immediately quenched in 4x Laemmli buffer containing 5% β-ME. Percent covalent adduct formation was calculated using Bio-Rad ImageLab software. The background subtraction function of ImageJ was used to process all gel images post-analysis.

### HUH-SEQ ssDNA LIBRARY CLEAVAGE, LIBRARY PREPARATION, AND SEQUENCING

A 90-nt ssDNA library with a central 7 base randomized region flanked by conserved regions harboring primer binding sites at either termini (7N ssDNA library) was constructed using IDT oPools service consisting of 128 individually synthesized DNA oligos mixed at equal molarity (SI Table). This method produced a minimally biased distribution of all 16,384 possible kmers (SI, histogram of oPools vs. ultramer). Recombinant Rep cleavage of the 7N ssDNA library was carried out in triplicate in 3 µM Rep and 300 nM (83.4 ng/µL) ssDNA library in 50 mM HEPES, 50 mM NaCl, and 1 mM MnCl2 for 1 hour at 37°C. The Rep enzymes were immediately heat inactivated by boiling at 95°C for 3 minutes. The remaining uncleaved ssDNA library from each Rep in-vitro cleavage reaction was diluted 10-fold in water and amplified using 0.5 µM TruGrade/HPLC purified primers from IDT containing Nextera adapters and spacer regions with 2x CloneAmp™ HiFi PCR Premix for 30 cycles. The resulting product was a 200 bp dsDNA amplicon run on a 1.5% agarose gel and stained with SybrSafe (SI, agarose gel image of one trial). Each 200 bp product was gel extracted (NucleoSpin Gel and PCR Clean-up kit, Machereny-Nagel) and eluted in 30 µL NE buffer resulting in samples of 30-60 ng/μL. All samples were barcoded with Illumia dual-indexing sequences via the Nextera adapters (University of Minnesota Genomics Core). Indexed samples are were pooled and run on a 1.5% agarose gel; the 270 bp barcoded pooled sample was gel extracted and then sequenced using a single Illumina HiSeq lane (350,000,000 paired-end reads, Genewiz) spiked with 30% PhiX.

### HUH-SEQ READ COUNT REDUCTION ANALYSIS AND SEQUENCE LOGO GENERATION

Raw NGS sequence data were processed using R. Non-randomized portions (e.g. adapter sequences) were removed from each read to extract only the randomized 7-mer (*k*-mer). 7-mers from reverse reads were reverse-complemented, and frequency counts for each of the 16,384 unique 7-mers were generated for the reference library from each of the Rep treatment libraries. Each treatment was then compared against the reference to estimate a log2-fold-change and percent reduction (reference – treatment/ reference) for each of its 7-mers. The percent reduction data was used to generate weighted sequence logos for each Rep using the ggseqlogo package in R. In addition, log counts per million (LogCPM), one-way ANOVA *F*-test statistics (F), p-values, and False Discovery Rate (FDR) statistics were generated using the edgeR package for each *k*-mer per Rep treatment in triplicate.

### EXTRACTING PREDICTED ORTHOGONAL REP HUH-TAGS AND *K*-MER SETS

Orthogonality of Reps was determined *in silico* using a custom R script. The script first iterates through each Rep and labels it as strongly reactive, moderately reactive, or nonreactive with each of the *k*-mers, with any log2FC under −3.0 considered strongly reactive, and any over −0.3 considered nonreactive. Then, the number of strongly-reactive-plus-nonreactive *k*-mers is counted for every possible pairing of Reps. Two Reps, A and B, are labeled as “likely orthogonal” if there exists at least one such *k*-mer in each direction—one where A is strongly reactive and B is nonreactive, and another where A is nonreactive and B is strongly reactive.

## Data Availability

Co-crystal structure coordinates and structure factors of PCV2^Y96F^ + 10-mer, WDV^Y106F^ + 10-mer, and WDV^Y106F^ + 8-mer complexes were deposited with accession codes 6WDZ, 6WE0, and 6WE1, respectively, in the Protein Data Base (PDB).

## Acknowledgements

This research was supported by an NIH NIGMS R35 GM119483 grant. EJA and RLE received salary support from Biotechnology NIH T32GM008347 and Muscle T32AR007612 Training Grants, respectively. HA received funding from NIH GM118047. WRG is a Pew Biomedical Scholar. This work is based upon research conducted at the Northeastern Collaborative Access Team beamlines, which are funded by the National Institute of General Medical Sciences from the National Institutes of Health (P30 GM124165). This research used resources of the Advanced Photon Source, a U.S. Department of Energy (DOE) Office of Science User Facility operated for the DOE Office of Science by Argonne National Laboratory under Contract No. DE-AC02-06CH11357.

## Author contributions

KJT, LAL, LP, LKL, and RLE grew protein crystals, processed x-ray data, and refined structures. KS collected data at the beam line. KJT and EJA conceived HUH-seq. KJT and MH developed and performed HUH-seq analysis. KJT designed biochemical experiments. KJT, LAL, AJN, LKL and BAE performed cloning, expressed and purified protein, and performed biochemical assays. KJT and WRG prepared and wrote the manuscript. All authors contributed to manuscript editing and gave approval of the final manuscript version.

## Supplementary Information

**Supp. Note 1: Scissile phosphate completes coordination of manganese in rep active sites**

The PCV2^Y96F^ + 10-mer active site contains an HUQ motif rather than an HUH motif along with a structurally conserved Glu48 that coordinates a manganese ion. Octahedral coordination of the ion is completed by an adjacent water molecule, which may be positioned by Arg54 or Glu100, and the O1 and O3’ of the scissile phosphate. A sequence conserved lysine, Lys99, may act as a general base and deprotonate the catalytic tyrosine poised near Phe96. The WDV^Y106F^ + 10-mer active site reveals identical coordination of manganese by the HUH motif and scissile phosphate. In contrast however, it uses Glu110 to coordinate the metal ion instead of the predicted Glu49 residue. This seems to result in shifting Lys109 out of position to act as a general base. It is unknown whether this coordination is a crystallographic artifact or if geminivirus Reps use a different cleavage mechanism that does not rely on a general base to activate the tyrosine. Rep and relaxase active sites are highly similar and perform the same general DNA cleavage mechanisms, using divalent metal ion coordination of O1 and O3’ of the scissile phosphate. This polarizes the partial positive charge of the phosphate catalyzing a nucleophilic attack by an adjacent active residue, though relaxases use a second conserved tyrosine for the DNA re-joining reaction ^1^.

**Supp. Fig. 1:**
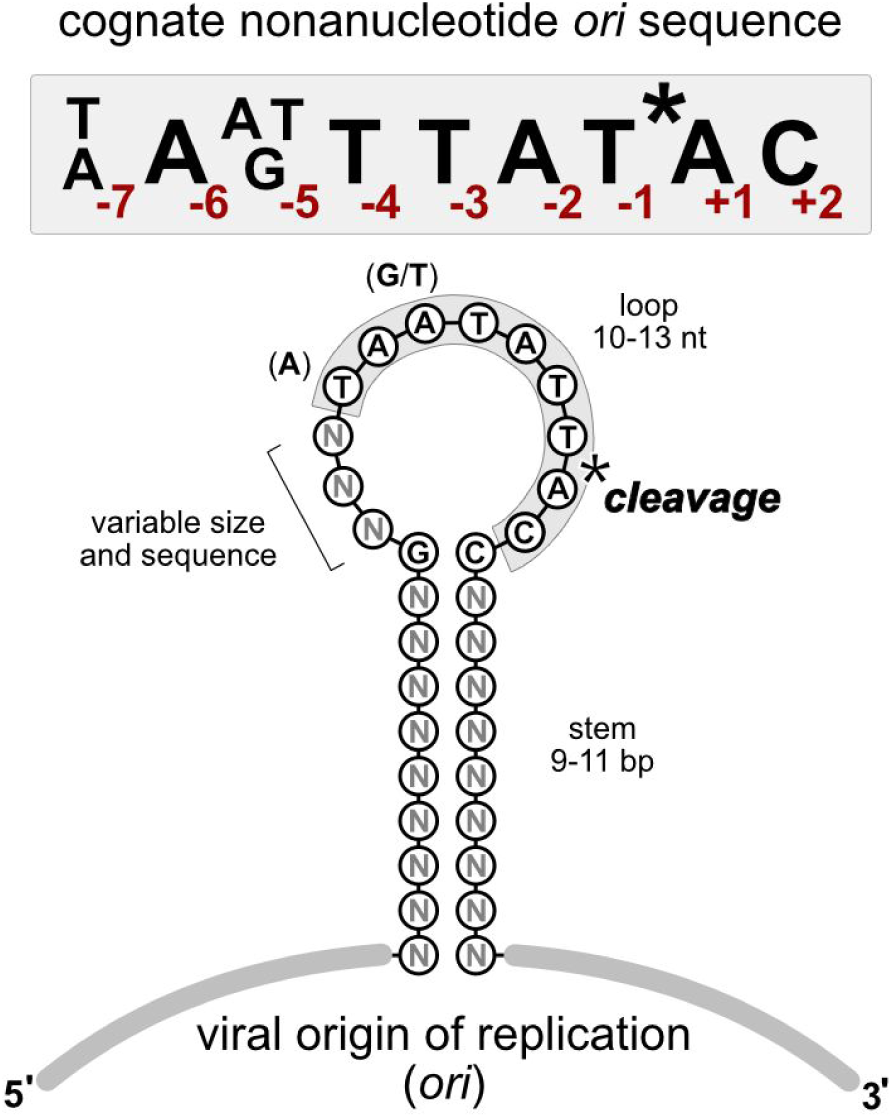
Hairpin structure and consensus cognate sequence recognized by panel of Reps. Consensus cognate nonanucleotide *ori* sequence of 10 different Reps from *Circoviridae, Nanoviridae, and Geminiviridae* (see Table 1 and SI). The origin or replication (*ori*) from these ssDNA viruses contains a stem-loop hairpin with Rep cleavage occurring between position −1 and +1 within the nonanucleotide sequence. The viral *ori* contains a stem that varies in sequence and between 9-11 base pairs in length while the loop contains the cognate nonanucleotide sequence and varies between 10-13 nucleotides in length.

**Supp. Fig. 2.**
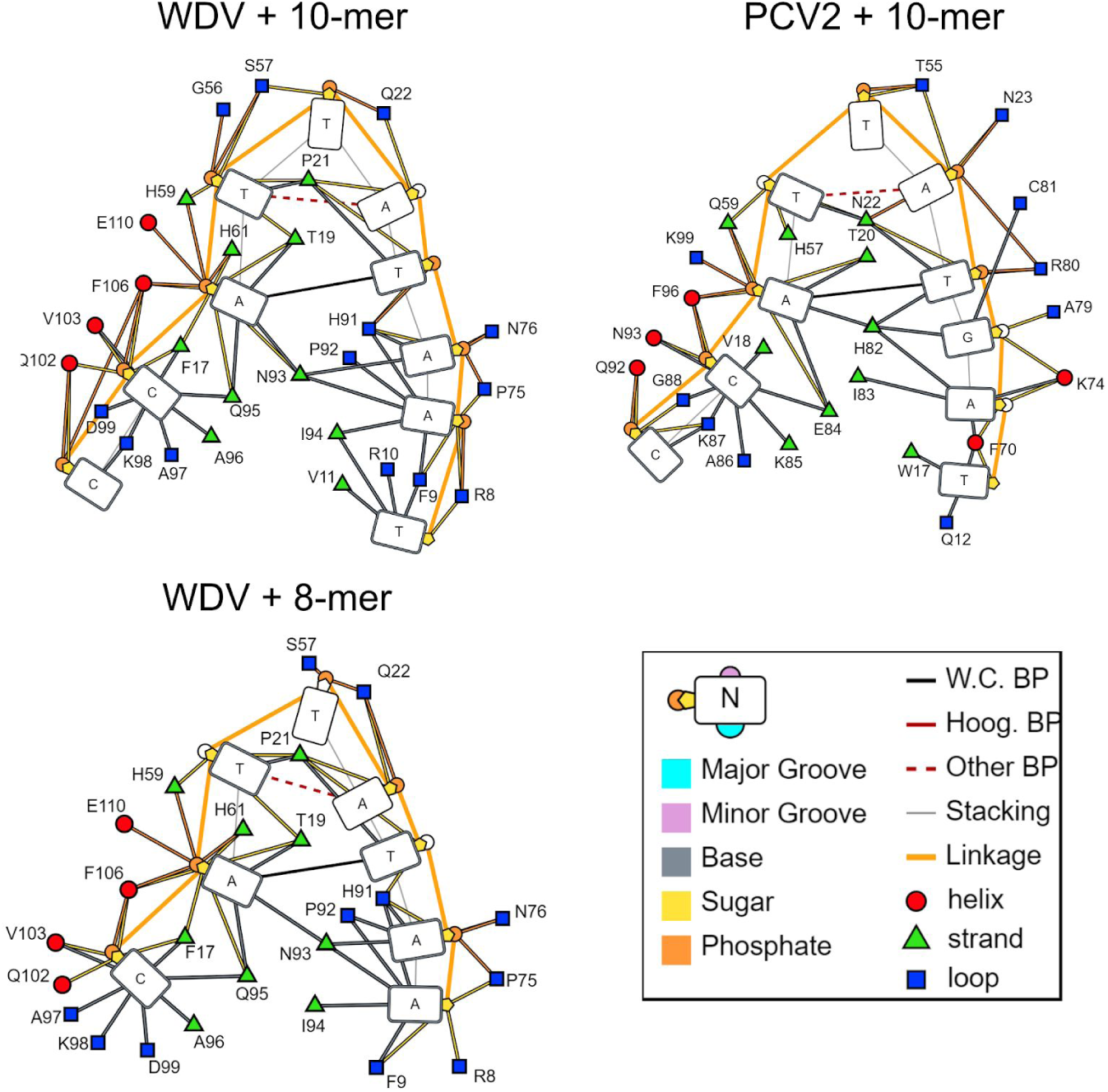
DNAproDB analysis of PCV2^Y96F^ and WDV^Y106F^ bound to a 10-mer and WDV^Y106F^ bound to 8-mer structures using default settings displaying contacts within 4 Å of the nucleotide backbone, sugar, and phosphate. Figure annotation and visualizations are taken directly from analysis software.

**Supp. Fig. 3:**
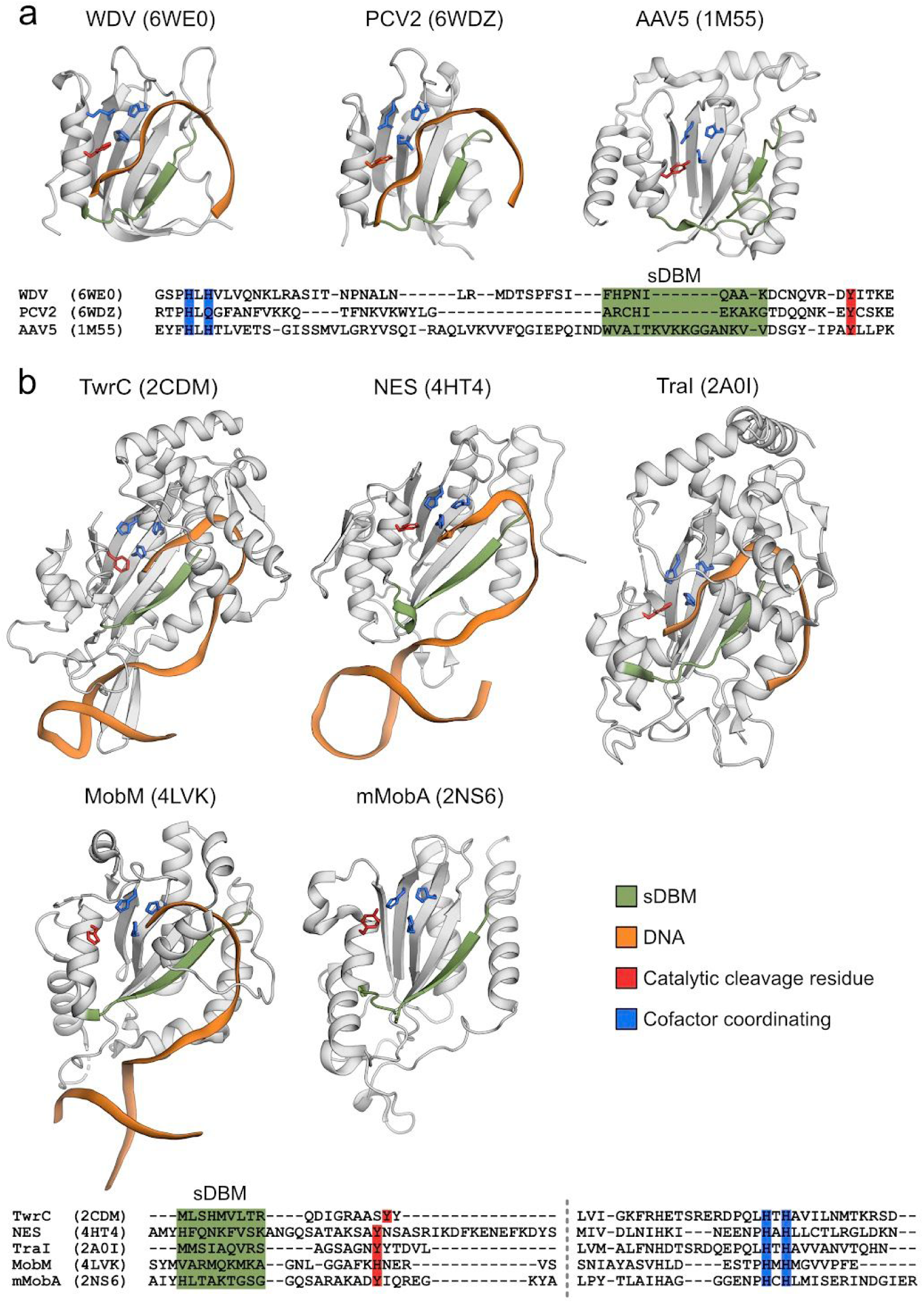
The sDBM is a conserved ssDNA binding motif spanning HUH endonuclease classes. Structural alignment using PROMALS3D and structural cartoons renderings highlight the relative positions of the sDBM (green) in **a**, *β*4 for Reps in and **b**, *β*1 for relaxase and transposases. Sections of the structural alignment consisting of the sDBM, the catalytic residue highlighted red, and the HUH-motif highlighted in blue are shown. Structures of protein-DNA complexes were used where available. DNA is shown as orange cartoon ribbon following C3 in ribose. Any bound metal ions are hidden.

**Supp. Note 2**

Sequential truncations of this geminivirus *ori* sequence of either the 5’ end or 3’ end indicated that at most, the nonameric sequence is necessary for sufficient cleavage activity because all ten Reps produced high adduct formation when the full nonameric sequence was retained in the target oligo. Perhaps the position −7 nucleotide is not necessary, because adduct formation was nearly identical in the absence of this nucleotide. Strikingly, CpCDV, and to a smaller extent PCV2 and CLCV, retain cleavage activity when only nucleotides in positions −2 through +2 are present (T_-2_T_-1_*A_+1_C_+2_). In contrast, TYLCV retained cleavage activity only when 8 of the 9 positions are retained in the target sequence, negating position −7 (A_-6_A_-5_T_-4_A_-3_T_-2_T_-1_*A_+1_C_+2_).

Single substitutions along the target sequence had no broad effect on the geminivirus Reps, except TYLCV activity was hampered by substitutions at positions −6 through +2. The nanovirus Reps, FBNYV and BBTV, have nearly identical target sequence profiles to that of TYLCV but tolerate both a transition and transversion at position −5. Interestingly, several double substitutions flanking the cleavage site of the target sequence were largely tolerated by geminivirus Reps, again with the exception of TYLCV, but cleavage of these sequences by circovirus and nanovirus Reps were abrogated.

**Supp. Fig. 4:**
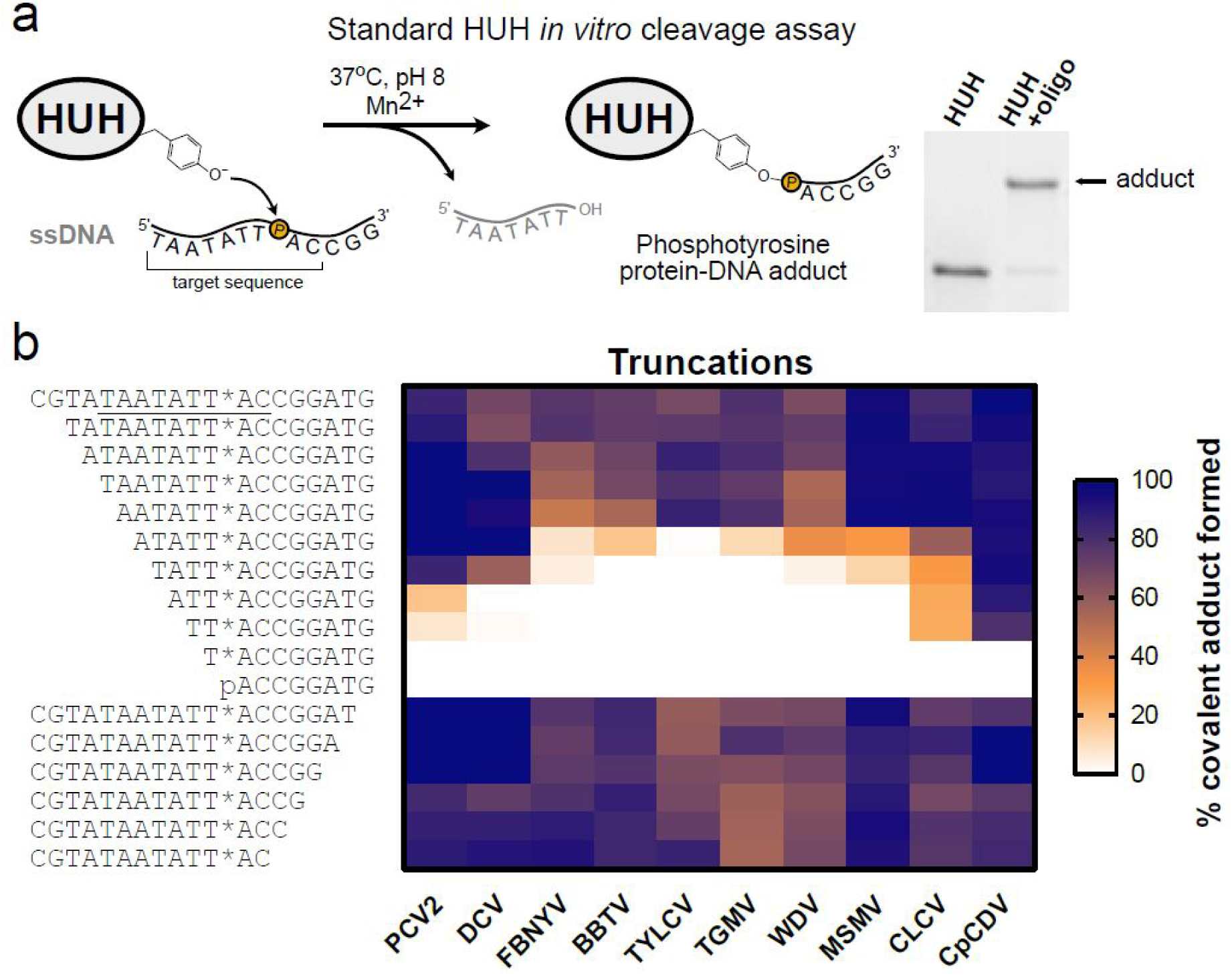
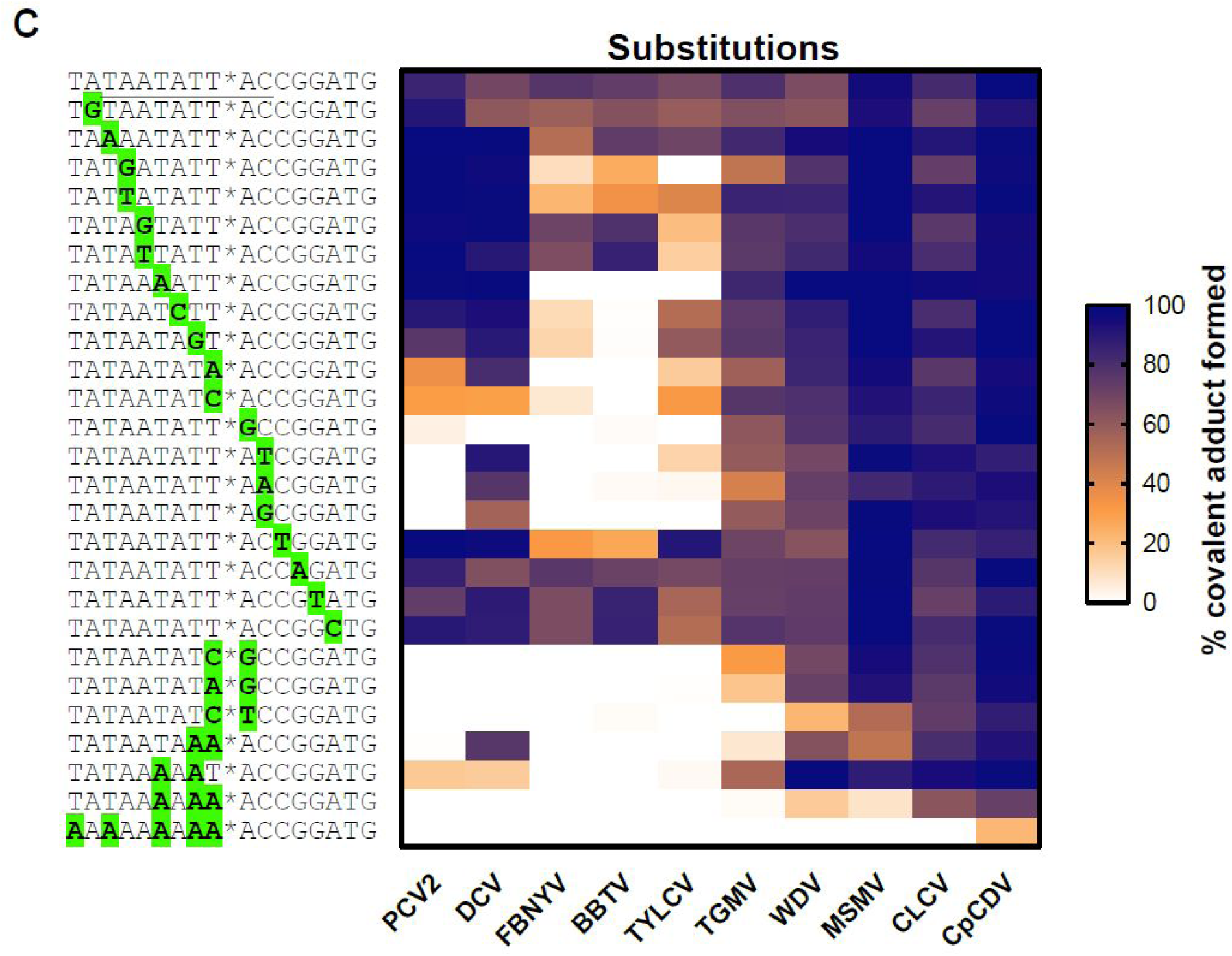
Standard *in vitro* cleavage assay with Rep panel and oligo library. **a**, The standard in vitro HUH reaction schematic where the catalytic tyrosine forms a covalent adduct with the phosphate at the +1 position scissile phosphate in the target sequence of a synthetic oligo. The phosphotyrosine adduct is stable under denaturing conditions evident by an upward shift from SDS-PAGE analysis. **b**, The truncations heatmap displays Rep covalent adduct formation with sequential truncations of synthetic DNA oligos harboring partial geminivirus *ori* target sequence with the cognate nonanucleotide *ori* sequence underlined. **c**,. The substitutions heatmap displays HUH-endonuclease covalent adduct formation with single or multiple substitutions within synthetic DNA oligos harboring the geminivirus *ori* target sequence, either 26nt or 17-nt in length (the entire 26-nt sequence is not shown in the figure). Full sequences of synthetic oligos used are provided. The conjugation reaction is carried out under the standard HUH in vitro reaction conditions with a 1:10 Rep:oligo ratio for 30 minutes. Each cell reflects a percent covalent adduct formation for a single reaction.

**Supp. Fig. 5:**
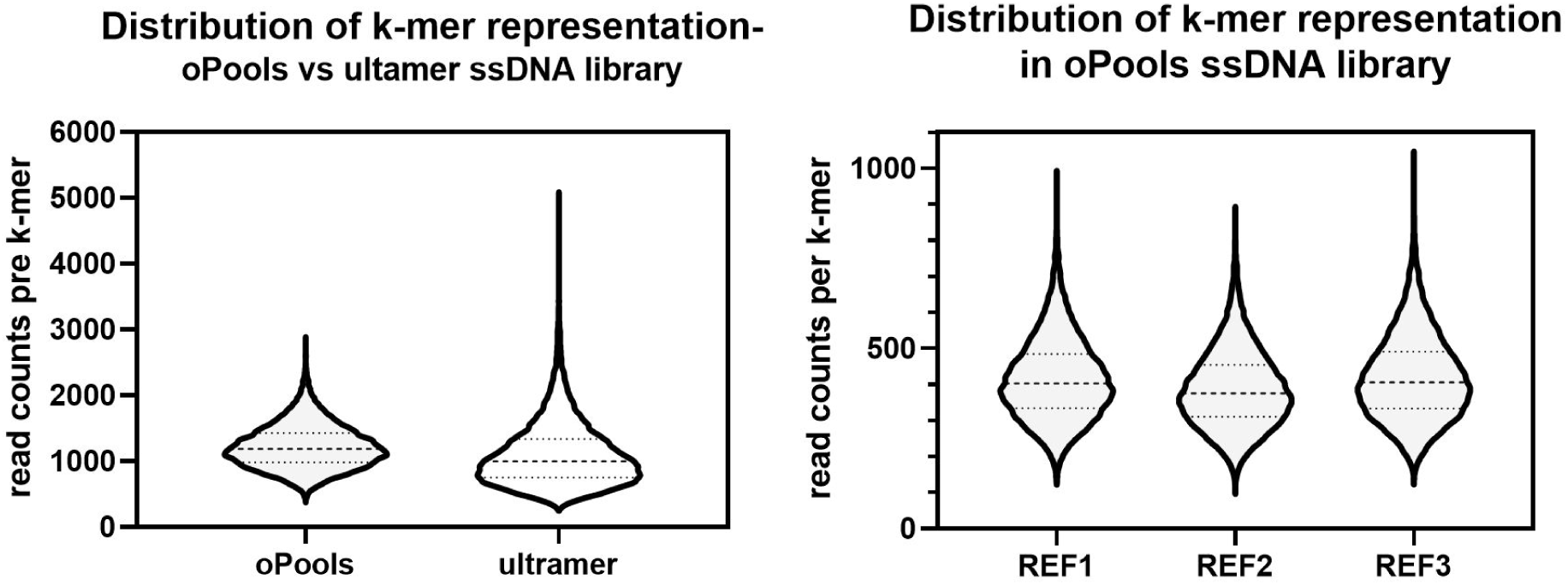
Distribution of *k*-mer representation in 7N ssDNA library for HUH-seq. Violin plot depicting the distribution of all 16,384 *k*-mers in the oPools generated 7N ssDNA library versus the ultramer generated 7N ssDNA library based on a sum of all read counts at each *k*-mer position from three untreated replicates. B: Violin plot depicting distribution of all 16,384 kmers present in each of the untreated reference (REF) replicates.

**Supp. Note 3**

Ideally, every sequence in the library must be at a high read count (>100 reads) since HUH-seq is a read count reduction based assay, so we limited the library size (SI Eq. 1) based on the total read counts provided by the HiSeq platform. We reasoned the size of the library, number of treatments, number of replicates, total read counts desired per *k*-mer, and percent of PhiX spike-in would all contribute to how large a library could be used (SI Eq. 2). We settled on creating a ssDNA library with 7 randomized nucleotides, which would produce on average 415 read counts per *k*-mer for 24 samples in triplicate (Eq. 1 and 2.).

**Supp. Equations 1 and 2:**

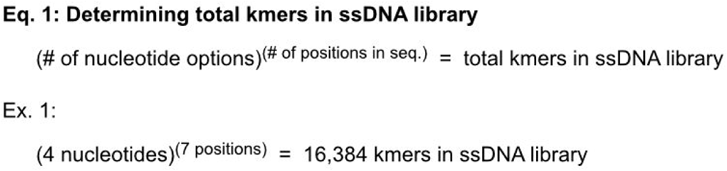

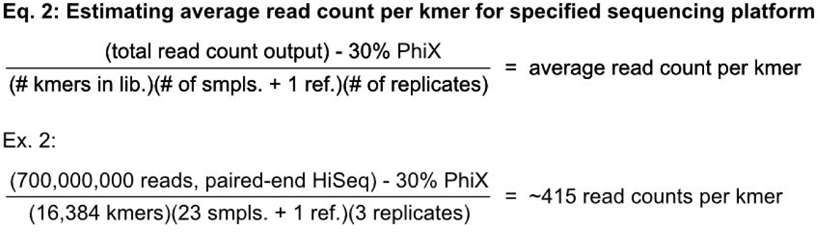

Eq. 1 determines the number of total kmers of user specified length in the synthetic ssDNA library dictated by the number of nucleotide options and number of randomized positions. Eq. 2 is used to estimate the average read count per kmer dictated by estimated total read count given by a NGS platform minus the recommended 30% PhiX spike in, all divided by the number of kmers in the ssDNA library, number of HUH-endonuclease samples and reference sample, and the number of replicates. This can be used to help determine which NGS platform will give sufficient read counts per kmer.

**Supp. Note 4: HUH-seq cleavage assay caveats and controls**

Lower concentrations of WDV had minimal impact on the ssDNA recognition profile of WDV, yet the lower maximum average percent reduction values correspond to fewer *k*-mer sequences being cleaved (Supp. Fig. 6a). Removing SUMO from WDV minimally affected the ssDNA recognition profiles as well (Supp FIg. 6b). Interestingly, the inactive WDV^Y106F^ mutant control revealed *k*-mers with significant percent reduction over the reference library but to a much lesser extent than wild-type WDV. We have verified that this inactive WDV mutant does not cleave the 26-nt geminivirus nonanucleotide *ori* sequence oligo using the standard in vitro cleavage assay (Supp Fig. 6c and 6d). We reasoned that the WDV^Y106F^ is still able to tightly bind the preferred target sequence and decrease the read count perhaps by partially blocking amplification of sequences bound to Rep rather than physical separation of the PBS sites due to cleavage. This result indicates that decreased read counts may be a consequence of both Rep binding and cleavage at a detectable yet much lower rate. Further, the ssDNA recognition profile generated for WDV^y106F^ is almost identical to that of WT WDV. We could refine this assay in the future with a more effective Rep inactivation step ensuring that cleavage is the only readout for the assay.

Another caveat we noticed is that the MSMV sequence logo did not show a prefered target sequence profile similar to that of the cognate nonanucleotide *ori* sequence. We presume that MSMV had a high cleavage rate in the constant region of the 7N ssDNA library somewhere near the 5’ end of the randomized region resulting in a largely random sequence logo (Supp. Fig. 6e). Optimizing the sequence of flanking constant regions of the 7N ssDNA library to limit cleavage in this region may allow us to generate an accurate target sequence cleavage profile of MSMV in the future. Simple optimization could eliminate these caveats, though it would give uncritical advantages.

Unknown binding, cleavage, and release kinetic factors may differ for each Rep and could be convoluting our ability to compare specificity. Ascertaining a full kinetic profile of each of these steps may give a better comparative picture of sequence specificity. In some cases, a high protein to substrate concentration ratio is known to negatively impact sequence specificity even of highly specific DNA-binding proteins such as zinc finger nucleases ^67^.

**Supp. Fig. 6:**
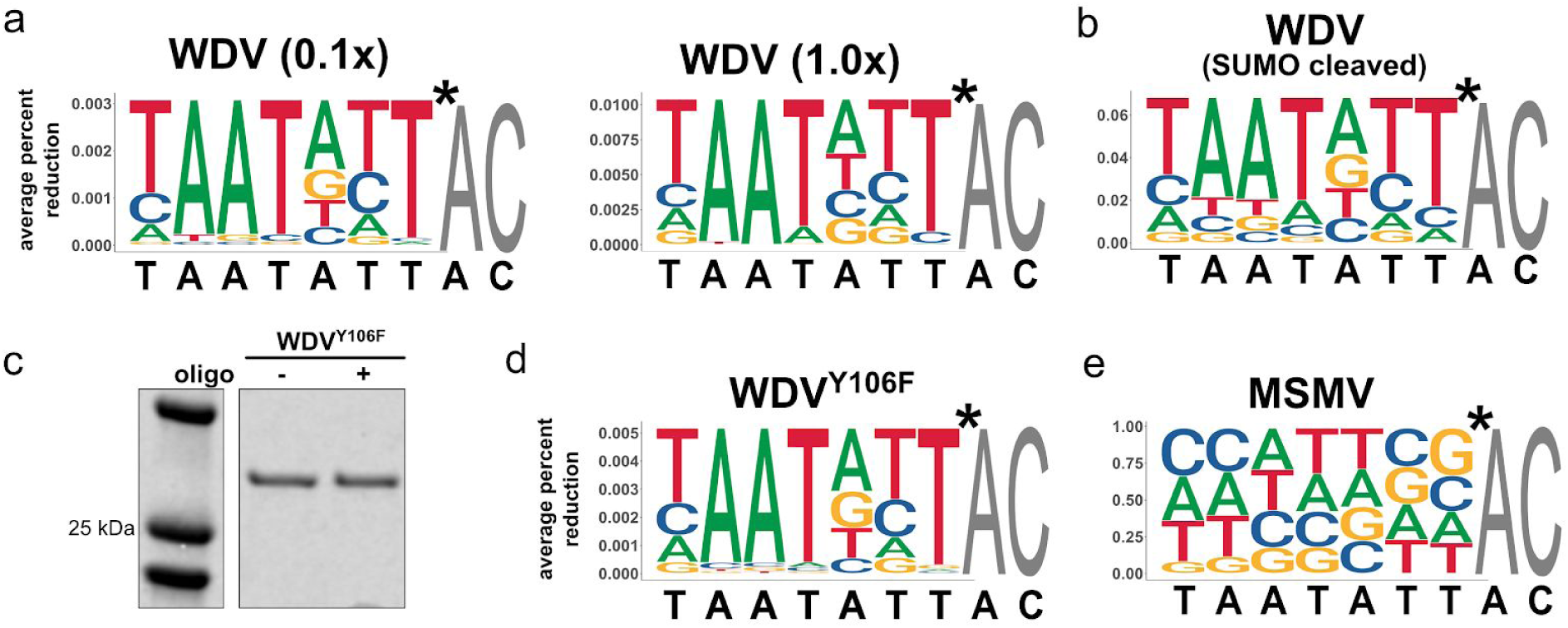
Effects of enzyme concentration dependence, tag-less enzyme, and inactive enzyme on HUH-seq readout. **a**, The weighted sequence logos of WDV at two lower concentrations, ten-fold less (x0.1) or equimolar (x1.0) WDV, in respect to the total 7N ssDNA library concentration. B. The weighted sequence logos of WDV with SUMO-cleaved. C. Recombinant SUMO-WDV^Y106F^ reacted with and without the 26-nt geminivirus *ori* sequence oligo under standard HUH in vitro assay conditions with 1:10 protein:oligo ratio. The SDS-PAGE gel scan is cropped to remove unrelated data. D. The weighted sequence logo WDV^Y106F^ and E. MSMV generated from HUH-seq analysis. Heights are scaled to represent the average percent reduction of each base at each position when compared to the reference library. Black sequences below each logo of the cognate nonanucleotide *ori* sequences from each respective virus.

**Supp. Table 1:**
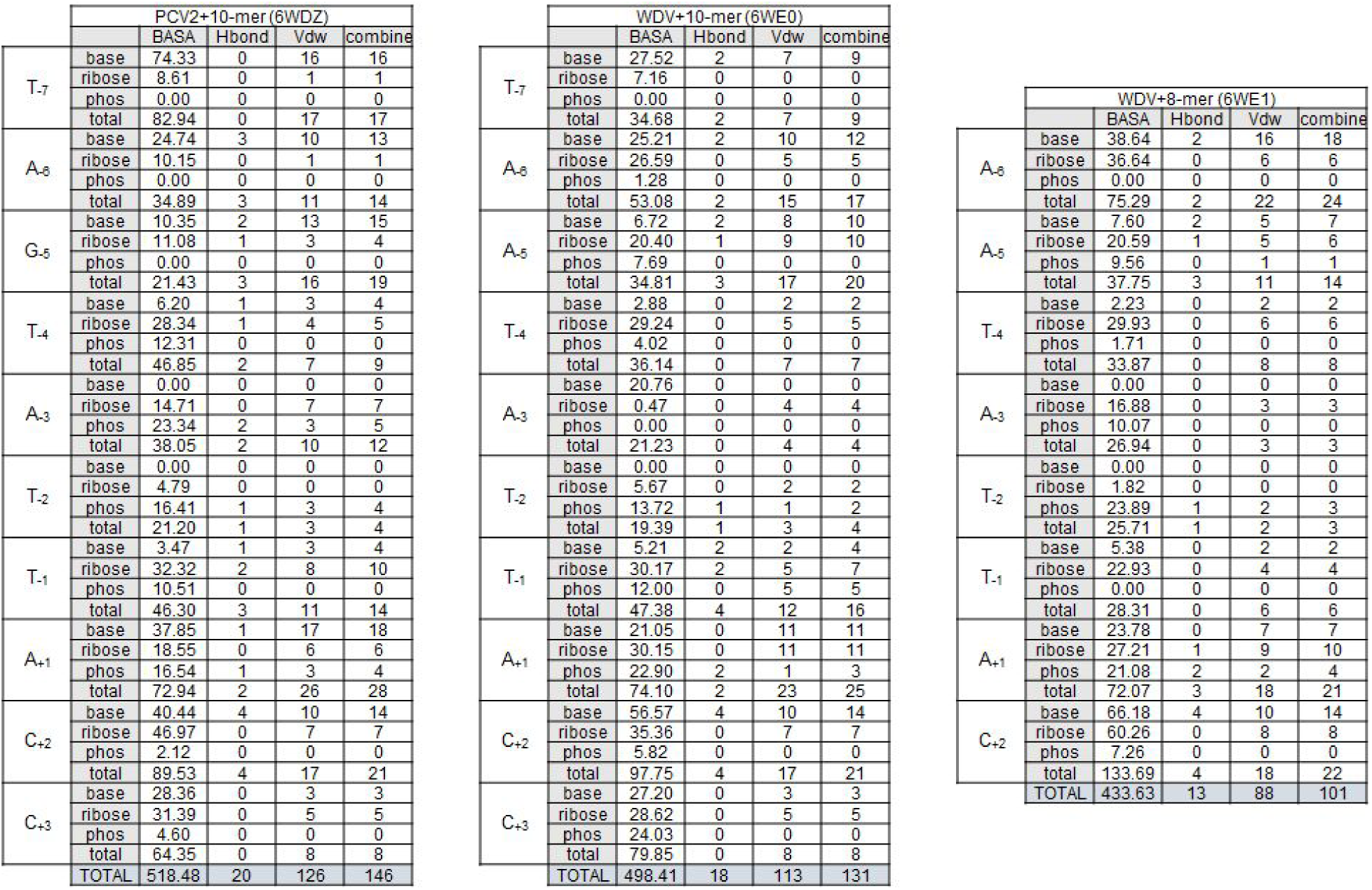
Individual BASA and contact values for protein-DNA interactions in co-crystal structures. Breakdown of all protein:DNA contacts and BASA values calculated by DNAproDB using default positions with ribose and phosphate interactions turned on.

**Supp. Figure 7:**
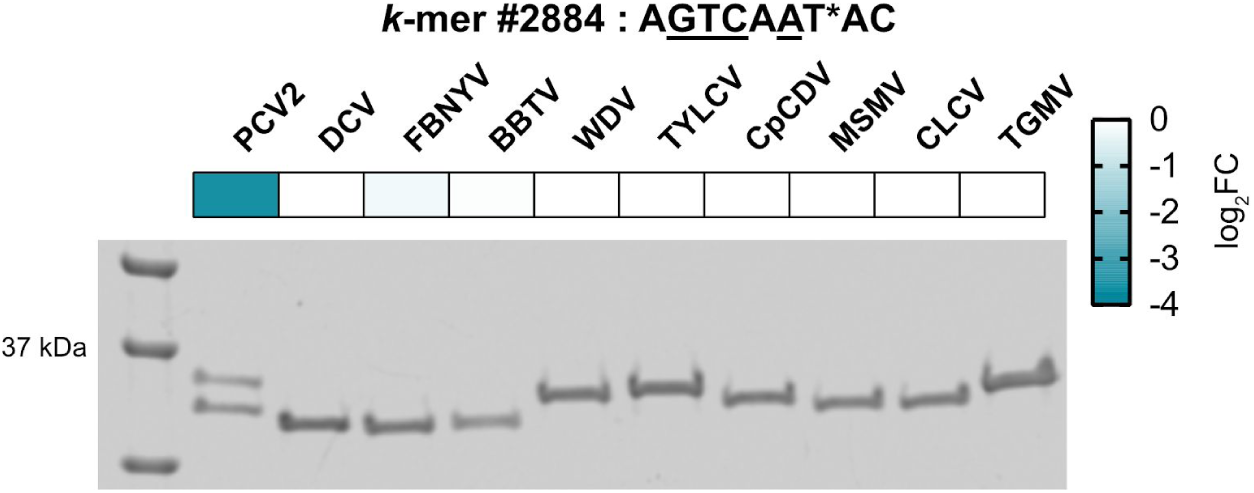
Discovery and validation of a target sequence exclusively cleaved by PCV2 Rep. Single row heat map shows log_2,_FC for each Rep treatment fromHUH-seq analysis for *k*-mer #2884, which bears a AGTCAAT sequence. There are four substitutions underlined at positions −6, −5, −4, and −2 with respect to the circovirus cognate nonanucleotide *ori* sequence. Below the heatmap is a respective gel highlighting covalent adduct formed under the standard HUH in vitro cleavage assay with a 1:2 protein:oligo ratio where only PCV2 forms adduct with *k*-mer #2884, and no detectable adduct is formed with any of the other 9 Reps.

**Supp. Table 2:**
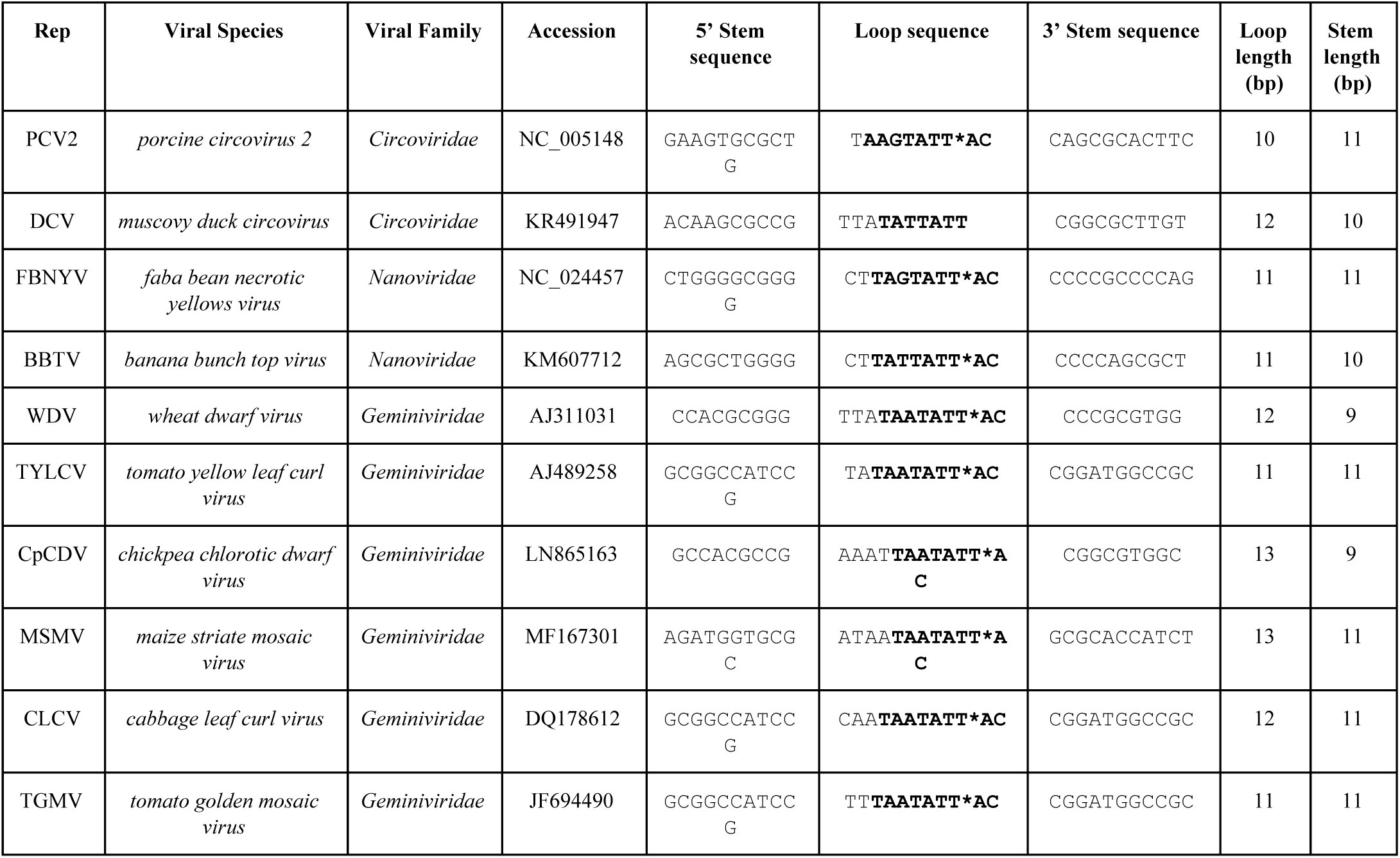
Panel of Reps.

**Rep protein sequences:**

>PCV2

PSKKNGRSGPQPHKRWVFTLNNPSEDERKKIRDLPISLFDYFIVGEEGNEEGRTPHLQGFANFVKKQTFNKVKWYLGARCHIEKAKGTDQQNKEYCSKEGNLLMECGAPRSQGQR

>DCV

MAKSGNYSYKRWVFTINNPTFEDYVHVLEFCTLDNCKFAIVGEEKGANGTPHLQGFLNLRSNARAAALEESLGGRAWLSRARGSDEDNEEYCAKESTYLRVGEPVSKGRSS

>FBNYV

MARQVICWCFTLNNPLSPLSLHDSMKYLVYQTEQGEAGNIHFQGYIEMKKRTSLAGMKKLIPGAHFEKRRGTQGEARAYSMKEDTRLEGPWEYGEFVP

>BBTV

MARYVVCWMFTINNPTTLPVMRDEIKYMVYQVERGQEGTRHVQGYVEMKRRSSLKQMRVFFPGAHLEKRKGSQEEARSYCMKEDTRIEGPFEFG

>WDV

MASSSTPRFRVYSKYLFLTYPQCTLEPQYALDSLRTLLNKYEPLYIAAVRELHEDGSPHLHVLVQNKLRASITNPNALNLRMDTSPFSIFHPNIQAAKDCNQVRDYITKEVDSDVNTAEWGTFVAVSTPGRKDRDAD

>TYLCV

MPRLFKIYAKNYFLTYPNCSLSKEEALSQLKKLETPTNKKYIKVCKELHENGEPHLHVLIQFEGKYQCKNQRFFDLVSPNRSAHFHPNIQAAKSSTDVKTYVEKDGNFIDFGVSQIDGRSARGGQQSANDAYAEAL

>CpCDV

MPSASKNFRLQSKYVFLTYPKCSSQRDDLFQFLWEKLTPFLIFFLGVASELHQDGTTHYHALLQLDKKPCIRDPSFFDFEGNHPNIQPARNSKQVLDYISKDGDIKTRGDFRDHKVSPRKSDAR

>MSMV

MSHTSFRFRAKNVFLTYPRCPIGPEFLCDHLWNLVTPYDPLYVHVAQENHKDGGLHSHVLIQTRIEISTFDPTYFDYTGTSIPGAVVFHPNIQACRNVRDCLAYIRKNTINEVSKGA

>CLCV

MPRNPKSFRLAARNIFLTYPQCDIPKDEALQMLQTLSWSVVKPTYIRVAREEHSDGFPHLHCLIQLSGKSNIKDARFFDITHPRRSANFHPNIQAAKDTNAVKNYITKDGDYCESG

>TGMV

MPSHPKRFQINAKNYFLTYPQCSLSKEESLSQLQALNTPINKKFIKICRELHEDGQPHLHVLIQFEGKYCCQNQRFFDLVSPTRSAHFHPNIQRAKSSSDVKTYIDKDGDTLVWGEFQVDGRSA

>PCV^Y96F^

PSKKNGRSGPQPHKRWVFTLNNPSEDERKKIRDLPISLFDYFIVGEEGNEEGRTPHLQGFANFVKKQTFN KVKWYLGARCHIEKAKGTDQQNKE**F**CSKEGNLLMECGAPRSQGQR

>WDV^Y106F^

MASSSTPRFRVYSKYLFLTYPQCTLEPQYALDSLRTLLNKYEPLYIAAVRELHEDGSPHLHVLVQNKLRAS ITNPNALNLRMDTSPFSIFHPNIQAAKDCNQVRD**F**ITKEVDSDVNTAEWGTFVAVSTPGRKDRDAD

>WDVc1

MASSSTPRFRVYSKYLFLTYPQCTLEPQYALDSLRTLLNKYEPLYIAAVRELHEDGSPHLHVLVQNKLRAS ITNPNALNLRMDTSPFSIF**RCHIE**AAKDCNQVRDYITKEVDSDVNTAEWGTFVAVSTPGRKDRDAD

